# Neurodevelopmental disorders and cancer networks share pathways; but differ in mechanisms, signaling strength, and outcome

**DOI:** 10.1101/2023.04.16.536718

**Authors:** Bengi Ruken Yavuz, M Kaan Arici, Habibe Cansu Demirel, Chung-Jung Tsai, Hyunbum Jang, Ruth Nussinov, Nurcan Tuncbag

## Abstract

Neurodevelopmental disorders (NDDs) and cancer are connected, with immunity as their common factor. Their clinical presentations differ; however, individuals with NDDs are more likely to acquire cancer. Schizophrenia patients have ∼50% increased risk; autistic individuals also face an increased cancer likelihood. NDDs are associated with specific brain cell types at specific locations, emerging at certain developmental time windows during brain evolution. Their related mutations are germline; cancer mutations are sporadic, emerging during life. At the same time, NDDs and cancer share proteins, pathways, and mutations. Here we ask exactly which features they share, and how despite their commonality, they differ in outcomes. Our pioneering bioinformatics exploration of the mutations, reconstructed disease-specific networks, pathways, and transcriptome profiles of autism spectrum disorder (ASD) and cancers, points to elevated signal strength in pathways related to proliferation in cancer, and differentiation in ASD. Signaling strength, not the activating mutation, is the key factor in deciding cancer versus NDDs.

## Introduction

NDDs arise from a dysfunctional nervous system during embryonic brain development. The origins of NDDs are still unclear. They may originate from dysregulation of neuron differentiation, during synapse formation and maturation, or other complex processes in the course of brain evolution, such as emergence from progenitor cells, neuron phenotypic specification, migration, and specific synaptic contacts. Flaws can result in faulty wired neuronal circuits (Nussinov et al., 2023, 2022a). Despite differing from processes associated with the emergence of cancer, data indicate that NDDs and cancer are related, with immunity likely the common factor. The immune and nervous systems coevolve as the embryo develops (Nussinov et al., 2022a). The outcomes, cancer or NDDs, reflect the different cell cycle consequences, proliferation in cancer and differentiation in NDDs. Proliferation requires a stronger signal to promote the cell cycle than differentiation does. This further suggests that in addition to nodes in the major signaling pathways, transcription factors (TFs) and chromatin remodelers, which govern chromatin organization, are agents in NDDs. Gene accessibility influences the lineage of specific brain cell types at specific embryonic development stages (Nussinov et al., 2023).

Here, we aim to uncover the shared features between neurodevelopmental disorders and cancer. We expect that these will help us understand the challenging question of how expression levels and mutations in the same pathways, and even the same proteins, including TFs and chromatin remodelers, can lead to NDDs versus cancer, with vastly different phenotypic presentations. Especially, we aim to discover what are the determining features deciding whether the major outcome is NDDs or cancer. We address this daunting goal by comprehensively leveraging mutations, transcriptomic data, and protein-protein interaction (PPI) networks. We compare the effects of mutations on the pathogenicity of commonly mutated genes in NDDs and cancer. We observe that mutations in NDDs tend to be weaker than those in cancer. To evaluate the pathway-level properties of NDDs and cancer, we reconstruct the disease-specific networks of autism spectrum disorder (ASD) and breast cancer and identify common TFs. Most of the targets of these common TFs are mutated in both ASD and breast cancer and involved in mitogen-activated protein kinase (MAPK), the cell cycle, and phosphoinositide 3-kinase/protein kinase B (PI3K/AKT) pathways. By using transcriptomic profiles of ASD and breast/brain/kidney cancers, we show that in breast cancer samples, there is an increase in signaling strength in shared pathways involved in proliferation and a decrease in differentiation. This, however, is not the case among ASD samples, where the signaling level is high in shared pathways involved in differentiation and low in proliferation.

Recent epidemiological studies on large cohorts of NDD patients demonstrated an increased risk for cancer compared to the general population. In one study, a standardized incidence ratio model was applied to a cohort of 8438 patients with autism retrieved from the Taiwan National Health Insurance database during 1997-2011. It implicated an increase in cancers of the genitourinary system and ovary among children and young adults (Chiang et al., 2015). Increased cancer risk was also observed in a population-based study among 2.3 million individuals with ASD from Nordic countries during 1987-2013 with co-occurring birth defects, including intellectual disability (Liu et al., 2022). A correlation between autism and cancer with shared risk factors was also pointed out (Kao et al., 2010). Another cohort study proposed that patients with bipolar disorder and their unaffected siblings have an especially higher risk of breast cancer compared to normal control groups (Chen et al., 2022). The association between brain, hepatocellular, and lung cancer among people with epilepsy was manifested by animal experiments, genotoxicity studies, and epidemiological observations. Possible underlying mechanisms have also been suggested (Singh et al., 2009, 2005).

NDD data has expanded recently, particularly *de novo* mutation data obtained by trio-sequencing and publicly available databases. However, it is still not as prevalent as the whole exome/genome sequencing data for cancer (Bragin et al., 2014; Turner et al., 2017). 32,991 *de novo* variants obtained from 23,098 trios are deposited in denovo-db (Turner et al., 2017). According to the database definition, *de novo* mutations are germline *de novo* variants present in children but not in their parents. The Deciphering Developmental Disorders (DDD) Study provides detailed genotype-phenotype information for 14,000 children with developmental disorders, and their parents from the UK and Ireland. Additionally, there are some knowledge databases with curated sets of genes and variants associated with one/multiple neurodevelopmental diseases or cancer (Abrahams et al., 2013; Piñero et al., 2015).

Here, we use *de novo* mutations in ∼10,000 samples with NDDs from denovo-db and somatic mutations of ∼10,000 tumor samples from The Cancer Genome Atlas (TCGA). Our large-scale analysis leads us to conclude that networks of NDDs and cancer can have shared proteins and pathways that differ in mechanisms, signaling strength, and outcomes. This conclusion is in line with our premise that cell-type specific protein expression levels of the mutant protein, and other proteins in the respective pathway and their regulators, the timing of the mutations, embryonic or sporadic during life, and the absolute number of molecules that the mutations activate, can determine the pathological phenotypes, cancer and (or) developmental disorders (Nussinov et al., 2022b). Our thesis is that these define the strengths of productive signaling (Nussinov et al., 2022c). In cancer, the major impact is on cell proliferation, while in NDDs it is on differentiation.

## Results

### NDD versus cancer mutations and networks data

NDDs and cancer are highly complex diseases caused by impairments in cellular processes such as cell growth, proliferation, and differentiation. This challenging complexity has led to the community’s desire to understand how their genetics, cellular environment, and signaling pathways are converging to express their distinct phenotypic outcomes (Jiang et al., 2022; Nussinov et al., 2022d; Parenti et al., 2020; Qi et al., 2016). Cancer results from gene alterations that provide cells with a growth advantage. Whereas numerous studies focused on the connection between the mutations—germline, *de novo*, or somatic—and cancer (Huang et al., 2018; Liu et al., 2020; Qing et al., 2020; Rashed et al., 2022; Stratton, 2008; Xu et al., 2020), the number of studies related to NDDs increased, though still lagging behind, far from reaching the same level. Qi et al. observed that among patients with NDDs, germline damaging *de novo* variants are more enriched in cancer driver genes than non-drivers (Qi et al., 2016). Bioinformatics analyses conducted on 219 cancer-related genes from Online Mendelian Inheritance in Man (OMIM, https://www.omim.org/about) and *de novo* mutations from 16,498 patients with NDDs, including ASD, congenital heart disease, and intellectual disability, found significantly more *de novo* mutations in cancer-related genes than in the 3391 controls (Li et al., 2020). In another study focusing on ASD, an evolutionary action method identified missense *de novo* variants that are most likely to contribute to the etiology of the disorder (Koire et al., 2021).

To identify genetic similarities and differences between NDDs and cancer, firstly we utilized publicly available mutation datasets. Public databases provide somatic mutation profiles of thousands of NDDs and tumor samples, including denovo-db and TCGA, respectively. Denovo-db includes *de novo* mutation profiles for 20 different NDD phenotypes for 9736 samples (Turner et al., 2017); TCGA covers 9703 samples with point mutations across 33 tissues (Figure 1A). Not all genes and their protein product variants affect the phenotypic output in the same way. Oncogenes, tumor suppressors, TFs, and chromatin remodelers are well-known examples of specific genes whose defects can cause observable alterations in phenotypic outcomes. We compared mutations and mutated proteins between *de novo* mutations in NDD data deposited in the denovo-db and TCGA, focusing on point mutations that affect only one residue in a protein. We identified 7908 genes in NDDs and 19,439 genes in TCGA with point mutations, among which 7838 genes are common. There are 147 oncogenes, 167 tumor suppressor genes, and 712 TFs in the NDD data, while 248 oncogenes, 259 tumor suppressor genes, and 1579 TFs are in TCGA. ∼40% of the mutated genes in TCGA also have mutations in NDD samples.

**Figure 1.**
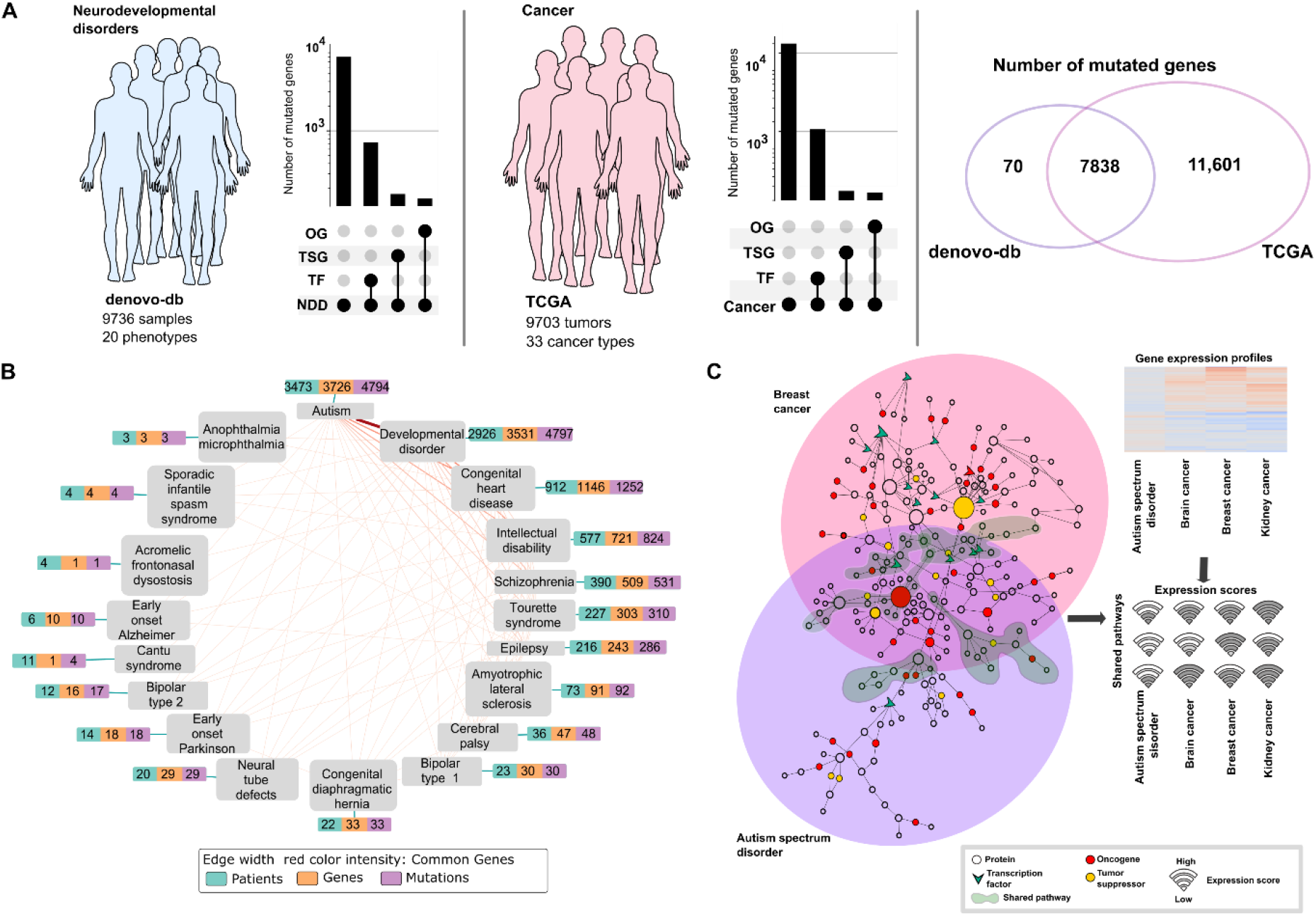
Overview of the data and workflow. **(A)** Statistics from NDDs and cancer datasets. Denovo-db deposits mutation profiles of 9736 samples with NDDs across 20 phenotypes (*left panel*). TCGA provides mutation profiles of 9703 tumors across 33 cancer types (*middle panel*). The length of each bar (*y*-axis in a logarithmic scale) in the upset plots shows the number of all mutated genes and the number of TFs, TSGs, OGs among the mutated genes for NDDs (*left panel*) and cancer samples (*middle panel*). There are 712 TFs, 162 TSGs, and 147 OGs out of 7907 mutated genes among NDD samples. Similarly, there are 1579 TFs, 259 TSGs, and 249 OGs out of 19,438 mutated genes among the cancer samples. The Venn diagram (*right panel*) shows that there are 7837 common mutated genes between NDDs and cancer; the number of NDD- and cancer-specific mutated genes are 70 and 11,601, respectively. Abbreviations: TSG, tumor suppressor gene; OG, oncogene. (**B**) Network of NDD phenotypes. Each node represents one phenotype in the network, and each edge represents the connection between two phenotypes if they share at least one commonly mutated gene. Each phenotype is represented with a vector of three numbers; the total number of patients having the phenotype (cyan), total number of genes carrying at least one mutation (orange), and total number of mutations associated with the phenotype (purple). The ticker edges represent the more commonly mutated genes. The most tightly connected pair among the phenotype pairs is autism and developmental disorder. (**C**) A conceptual representation of network comparison analysis between NDDs and cancer. Two distinct networks (*left panel*) reconstructed for breast cancer (large pink circle) and ASD (large purple circle). These two networks have both shared (shaded green) and separated regions. These networks contain oncogenes (red circle), tumor suppressors (yellow circle), and TFs (green chevron). The transcriptome analysis (*upper-right panel*) associates the expression levels of the nodes with the pathway activity. Each enriched pathway in the network can be quantified with the average expression level of its nodes, which is called “pathway scoring.” The score of each shared pathway (1, 2, .., *n*) for each disease (ASD, purple; cancer, red) is calculated (shown as a wifi icon where the higher score is the stronger signal).

The network of NDD phenotypes in the denovo-db database covers 20 NDD phenotypes with a varying number of patients, mutated genes, and mutations (Figure 1B). Only two of these phenotypes—autism and developmental disorders—have more than 1000 samples. In autism, there are 3473 patients, 3726 mutated genes, and 4794 mutations; in the 2926 samples of developmental disorders, there are 3531 mutated genes with 4797 mutations. In the network, the width of edges between the phenotypes is commensurate with the number of commonly mutated genes; autism and developmental disorders share the most. Congenital heart disease and intellectual disability have less than 1000 samples, 912 and 577, respectively. The remaining 15 phenotypes, including schizophrenia, epilepsy, and cerebral palsy, have less than 500 samples.

### Conceptualization, construction and comparisons of the networks, expression profiles, and mutation frequencies, in NDDs versus cancer

Figure 1C conceptualizes our study as follows: First, we reconstructed PPI networks using mutated genes in breast cancer and ASD as seeds. The networks we obtained include disease-specific regions as well as shared subnetworks for ASD and cancer. Then, we compared the expression scores of the pathways in the shared subnetwork by using gene expression profiles.

Our premise is that NDD mutations offer modest but prolonged signaling, whereas cancer mutations are associated with high signaling levels (Nussinov et al., 2022a, 2022b, 2022c, 2022d). Driver mutations are frequent, which is why they are often identified as drivers unless there is experimental data for potent rare mutations (Nussinov et al., 2019b; Nussinov and Tsai, 2015). Weaker or moderate mutations occur less frequently; otherwise, they are drivers. Similarly, the difference between passenger and driver mutations is also based on statistics; their counts are low. As one indicator of mutation strength, we compared the frequency of the cancer driver mutations in TCGA and NDD mutations amongst TCGA samples. For cancer driver mutations, we used the Catalog of Validated Oncogenic Mutations from the Cancer Genome Interpreter (CGI) (Muiños et al., 2021). Only missense or nonsense mutations were included in the analyses, which comprised 3688 driver mutations in 237 genes. Among 14,133 unique NDD mutations, 1504 are in TCGA (Figure 2A). On the other hand, TCGA harbors 1060 unique driver mutations. Interestingly, only 23 mutations are shared across known cancer driver mutations and NDDs (see the inset Venn diagram of Figure 2A). This finding suggests that although there are shared mutations between the two pathologies, these mutations tend to be on the weaker side in terms of a driver effect. In addition, compared to driver mutations, the mutations present in both NDDs and TCGA are notably rare in the TCGA cohort, as demonstrated by the difference in the mutation frequency distribution in TCGA with a *t*-test (*p* = 0.001). Therefore, when we limit the mutations to those present in TCGA, only ∼1% of NDD mutations are cancer drivers, and they have very low frequencies among TCGA samples. Figure 2B depicts the number of mutated samples in commonly mutated genes among NDDs and cancer. Most commonly mutated genes have more mutation hits at different positions among all cancer samples. Our observations point to only relatively few common NDDs and cancer driver mutations, making it crucial—even if difficult—to understand the mechanisms through which these common mutations impact gene function and disease phenotypes. We used pathogenicity scores from MutPred2 (Pejaver et al., 2020), which probabilistically predict the impact of variants on protein structure and function. We anticipate that variants may have an impact on protein structure, which can either stabilize or destabilize the conformation of the protein depending on protein function and disease phenotypes. The more harmful a mutation is, the closer its pathogenicity score is to one. A comparison of the distribution of the pathogenicity scores of the NDDs and driver mutations calculated using MutPred2 demonstrates that driver mutations have higher pathogenicity than NDD mutations (*t*-test, *p* < 5 × 10^-30^) (Figure 2C). We observe that most driver mutations accumulate in regions where the pathogenicity scores are larger than 0.8 on the *y*-axis. NDDs harbor mutations in key cancer genes such as *PTEN*, *PIK3CA*, *MTOR*, *KIT*, etc. These mutations have lower frequencies among tumor samples from TCGA, which is an indicator of the lower potency of these mutations. The number of residues hit by mutations among NDD samples is usually lower.

**Figure 2.**
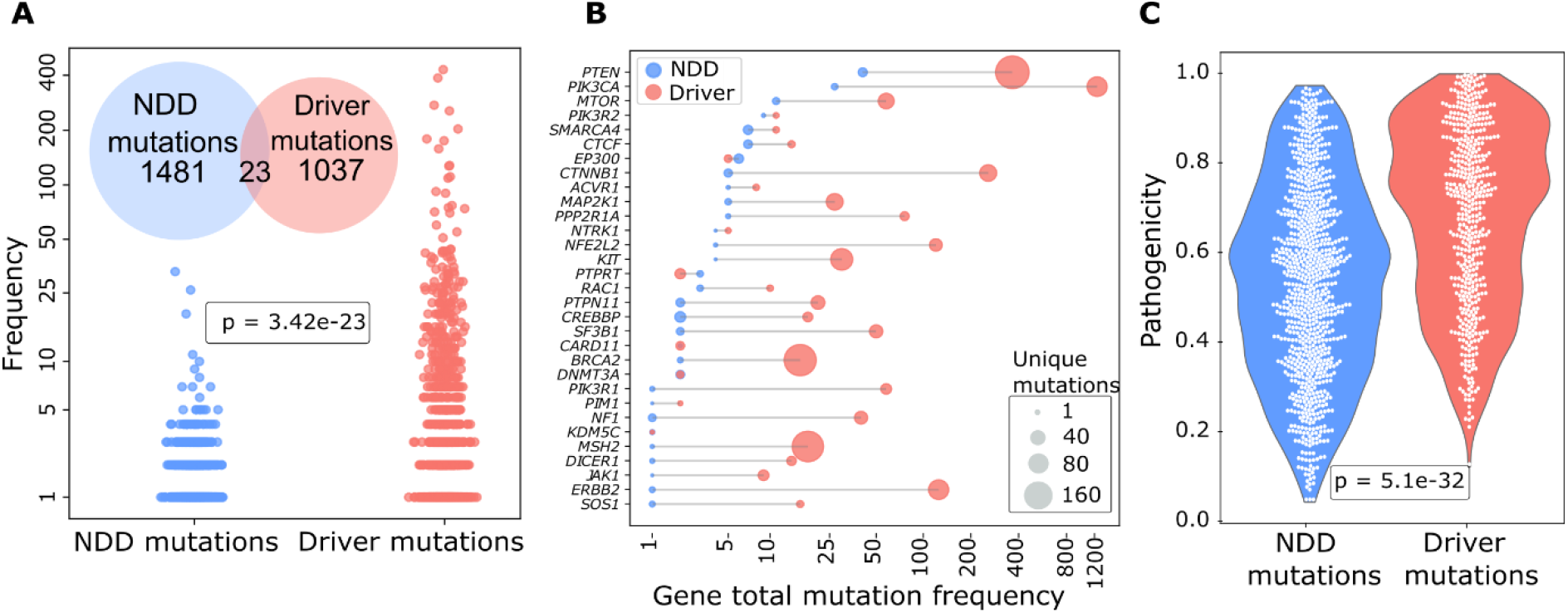
Comparison of mutations between NDDs and cancer. **(A)** Frequency-based analysis of mutations for NDDs and cancer. The cancer driver mutations in TCGA in comparison to the frequency of NDD mutations. The cancer driver mutations were selected amongst tumor samples only. Among the cancer mutations in TCGA, 23 mutations are shared between NDD and known cancer driver mutations, while 1481 are NDD-specific and 1037 are cancer-specific mutations (inset Venn diagram). Comparison of the frequency of these mutations in the TCGA cohort (*y*-axis in a logarithmic scale, where *frequency = log10(N+1)* and *N* is the number of patients). The difference between mutation frequency distribution in TCGA with *t*-test shows that the mutations present in both NDDs and TCGA are significantly rare in the TCGA cohort when compared to driver mutations (*p* < 0.001). (**B**) Frequency of mutations on common genes in NDDs and known cancer drivers datasets. The dumbbell plot shows the mutation frequencies of common genes–the genes harboring at least one point mutation among NDDs and cancer samples–in cancer (TCGA) and NDDs (denovo-db) simultaneously. Cancer driver mutations (red) are more frequent than or equal to NDD mutations (blue) except *EP300* and *PTPRT*. The size of the circles represents the number of unique mutations each gene carries. The *x*-axis in a logarithmic scale represents the number of patients having at least one mutation in the corresponding gene in TCGA or NDD sets. (**C**) MutPred2 pathogenicity scores of NDDs and cancer driver mutations. Violin plots show the distribution of NDD and driver mutation pathogenicity scores. A comparison of the pathogenicity scores using a *t*-test shows that the pathogenicity of driver mutations is significantly higher (*p* < 0.001). Pathogenicity scores are between 0 and 1, where 1 is the most pathogenic.

### Distribution of the locations of NDD and cancer mutations and modes of action

Phosphatase and tensin homolog (PTEN) phosphatase and PI3Kα lipid kinase are respectively negative and positive regulators in the PI3Kα/AKT/mTOR pathway. PTEN dephosphorylates phosphatidylinositol 3,4,5-trisphosphate (PIP_3_) to phosphatidylinositol 4,5-bisphosphate (PIP_2_) produced by PI3K. The signaling lipid PIP_3_ recruits AKT and PDK1 (phosphoinositide-dependent kinase 1) protein kinases to the plasma membrane, thereby playing a vital role in cell growth, survival, and migration (Jang et al., 2021; Zhang et al., 2019). Loss of function of PTEN by germline or somatic mutations leads to increased PIP_3_ concentrations at the membrane and promotes cell proliferation mediated by PI3Kα. Since the PI3Kα/AKT/mTOR pathway is one of the primary regulators of cell proliferation and differentiation, the mechanistic hallmarks of the mutations are vital to understand. Analysis of mutations in PTEN (Figure 3A) and PI3Kα (Figure 3B) sequences reveals that NDD mutations on these proteins usually occur at less frequently mutated sites among tumors (see Materials and Methods). R130* mutation in NDDs on PTEN is an exception, yet it is less frequent compared to the R130Q and R130G mutations at the same position in cancer.

**Figure 3.**
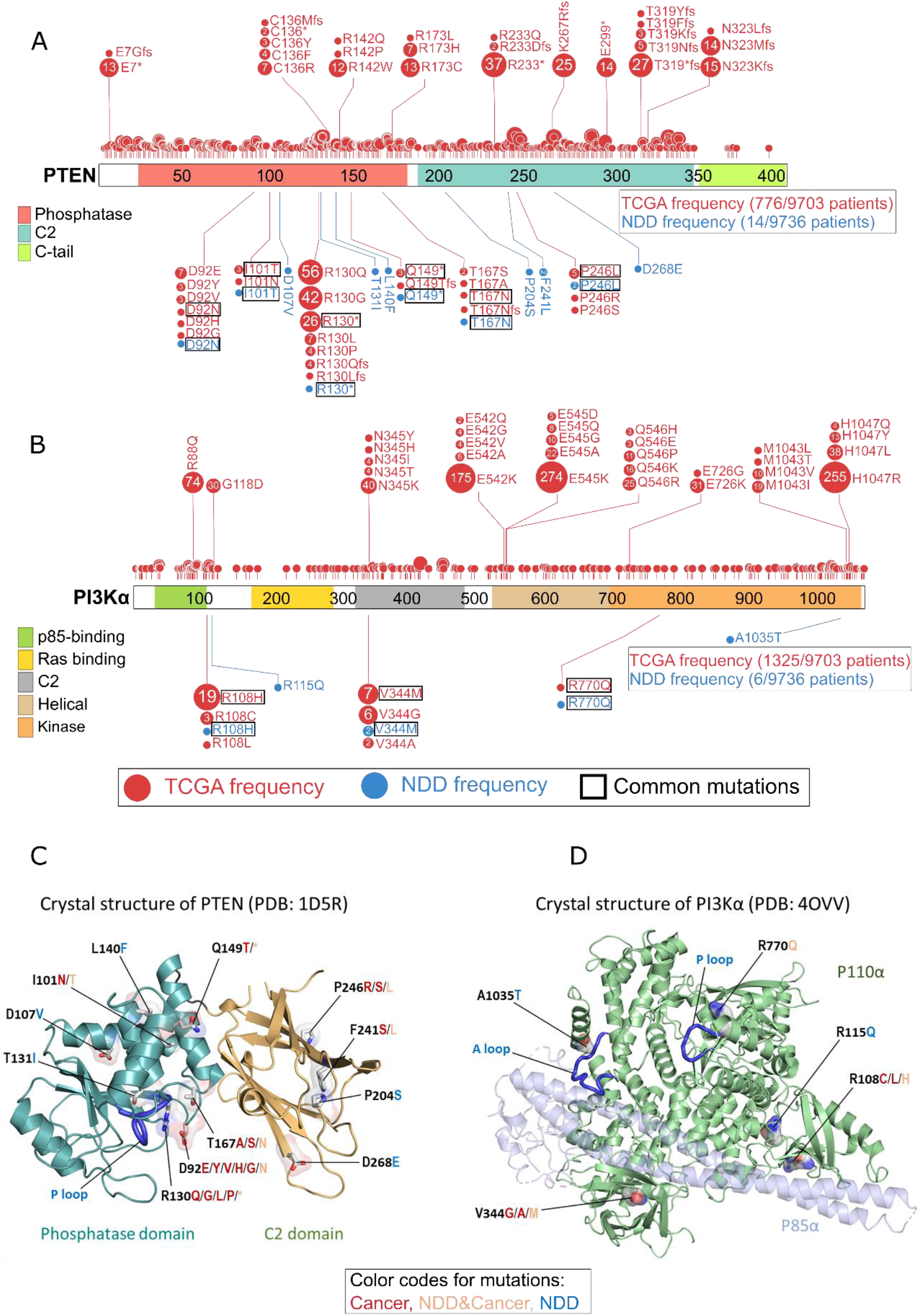
Profiles of TCGA and NDD mutations for PTEN and PI3Kα at the residue level on the sequence and structure. **(A)** Mutations of PTEN are shown as circles, where the phosphatase domain (red), C2 domain (dark green), and C-tail (light green) are represented as colored boxes along the sequence. The number and size of the circle represent the frequency of each mutation in the NDD (blue) or TCGA (red) datasets. Mutations shared by both datasets are highlighted with rectangular borders for emphasis. Total mutation frequencies and the total number of patients in each dataset are shown in the bottom right box. Nonsense mutations are abbreviated with star (*) sign. 6 of 12 PTEN mutations in the NDD set are present in TCGA. Only R130* has a high frequency relative to other shared mutations, yet it is much less frequent when compared to two other TCGA mutations on the same position, R130Q and R130G. (**B**) Mutations of PI3Kα (*PIK3CA*) are shown as circles where ABD (green), RBD (yellow), C2 domain (gray), helical domain (light orange), and kinase domain (orange) are represented as colored boxes along the sequence. The number and size of the circle represent the frequency of each mutation in the NDD (blue) or TCGA (red) datasets. Mutations shared by both datasets are highlighted with rectangular borders for emphasis. Total mutation frequencies and the total number of patients in each dataset are shown in the bottom right box. 3 of 5 PI3Kα mutations in the NDD set are present in TCGA. None of these TCGA mutations are on the most frequently mutated residues or among the most frequent mutations. Abbreviations: ABD, adaptor-binding domain; RBD, Ras-binding domain. The 3D structures of (**C**) PTEN (PDB: 1D5R) and (**D**) PI3Kα (PDB: 4OVV) with selected NDD and TCGA mutations. For each residue, mutated amino acids are colored in red, blue, or orange if they are present only among cancer, NDDs or both phenotypes, respectively. In PTEN, these mutations are known to affect the functions of protein including loss of phosphatase activity, reduced protein stability at the membrane, and failing to suppress AKT phosphorylation. In PI3Kα, these mutations may interrupt protein activation and reduce protein stability at the membrane.

While several residues of PTEN were mutated in both NDDs and cancer, some mutations—such as T131I, L140F, and D268E—are NDD-specific (Figure 3A). As to the domain distribution, among the NDD samples, mutated residues D92, I101, R130, T131, L140, Q149, and T167 are on the phosphatase domain, and P204, F241, P246, and D268 are on the C2 domain (Figure 3C). PTEN’s catalytic activity occurs in the phosphatase domain that contains the P loop (residues 123–130) with the catalytic signature motif, _123_HCxxGxxR_130_ (where x is any amino acid). PTEN mutations in the P loop, or nearby, such as at the residues R130 and T131, can directly constrain the P loop, leading to silencing PTEN catalytic activity. The mutation at residue D92 in the WPD loop (residues 88–98) can disrupt the closed WPD loop conformation that can bring D92 to the active site. D92 is involved in the catalytic activity during the process of hydrolysis to release the phosphate group from Cys124 after transferring it from PIP_3_. Other PTEN mutations, which are distant from the active site, can allosterically bias the P loop dynamics, reducing protein stability and its catalytic activity. A similar pattern is observed in PI3Kα; the rare mutations R108H, V344M, and R770Q are harbored in both NDDs and cancer, while R115Q and A1035T are specific to NDD samples (Figure 3B). V344 is on the C2 domain; R770 and A1035 are on the N- and C-lobes of the kinase domain, respectively (Figure 3D). R770 is located near the P loop, and R108 is on the interface of the catalytic subunit p110α and the regulatory subunit p85α. The mutations at these positions in PI3Kα may promote protein activation and increase protein stability at the membrane, but their mutational effects appear to be weaker than the driver mutations.

Several studies investigated germline mutations in PTEN and their association with tumor susceptibility or developmental disorders (Mighell et al., 2018; Portelli et al., 2021; Spinelli et al., 2015; Wong et al., 2018). For example, the rare I101T mutation on PTEN is present in NDDs and cancer samples. This mutation is identified as related to reduced lipid phosphatase activity and protein stability in a study conducted among 13 patients with PTEN hamartoma tumor syndrome (PHTS) who have autistic features, neurodevelopmental delays, and macrocephaly. The I101T mutant retained almost 30% of the lipid phosphatase activity of the wild-type protein; hence, it might be one of the major causes of tissue overgrowth and autistic appearance (Wong et al., 2018). Although available data are limited, PTEN retains its tumor suppressive function in NDDs while becoming fully dysfunctional in cancer samples.

### NDD- and cancer-specific networks regulate common pathways with different signaling outcomes

Although alterations in the same pathways and proteins contribute to the emergence of NDDs and cancer with different weights, the timing of the mutations, the number of activated molecules, the expression level of the mutated protein, and the proteins in the corresponding pathway have a major impact on the phenotypic outcome (Li et al., 2020; Nussinov et al., 2022d). To understand the divergence between these two pathologies, we analyzed NDD- and cancer-specific networks and compared the signaling outcomes of the pathways using gene expression values. We reconstructed ASD- and breast cancer-specific networks based on frequent mutations, comprising 168 driver genes in breast cancer, and 190 mutated genes that are present in at least three ASD patients. We extracted the graphlet motifs, small significant subnetworks, from the reference interactome HIPPIE through mutations with an unsupervised learning approach (Alanis-Lobato et al., 2017; Milenković and Pržulj, 2008). To select the most relevant interactions in a disease from the graphlet motifs with the PageRankFlux algorithm, we constructed the ASD-specific network with 350 proteins and 1291 interactions, and the breast cancer-specific network with 284 proteins and 1878 interactions (Supplementary Data 1) (Figure 4A) (Arici and Tuncbag, 2023; Rubel and Ritz, 2020). As can be expected based on our relatively weak mutation outcome premise of NDDs, some critical TFs such as Myc, p53, and Jun with cancer driver mutations are not frequently mutated in ASD. However, mutated genes can indirectly regulate these TFs in the ASD-specific network due to the rewiring of the signaling network. We found 23 common TFs in ASD- and breast cancer-specific networks. TF complexes including Myc/Max or Jun/Fos (AP-1, activator protein 1) regulate the expression of numerous target genes downstream the MAPK phosphorylation cascade in signal transduction (Garces de Los Fayos Alonso et al., 2018; García-Gutiérrez et al., 2019). Complexes composed of common TFs are primarily involved in cell cycle regulation through their targets, such as E2F mediating cyclin-dependent kinases (CDKs) in cell proliferation (DeGregori et al., 1997; Tadesse et al., 2019).

**Figure 4.**
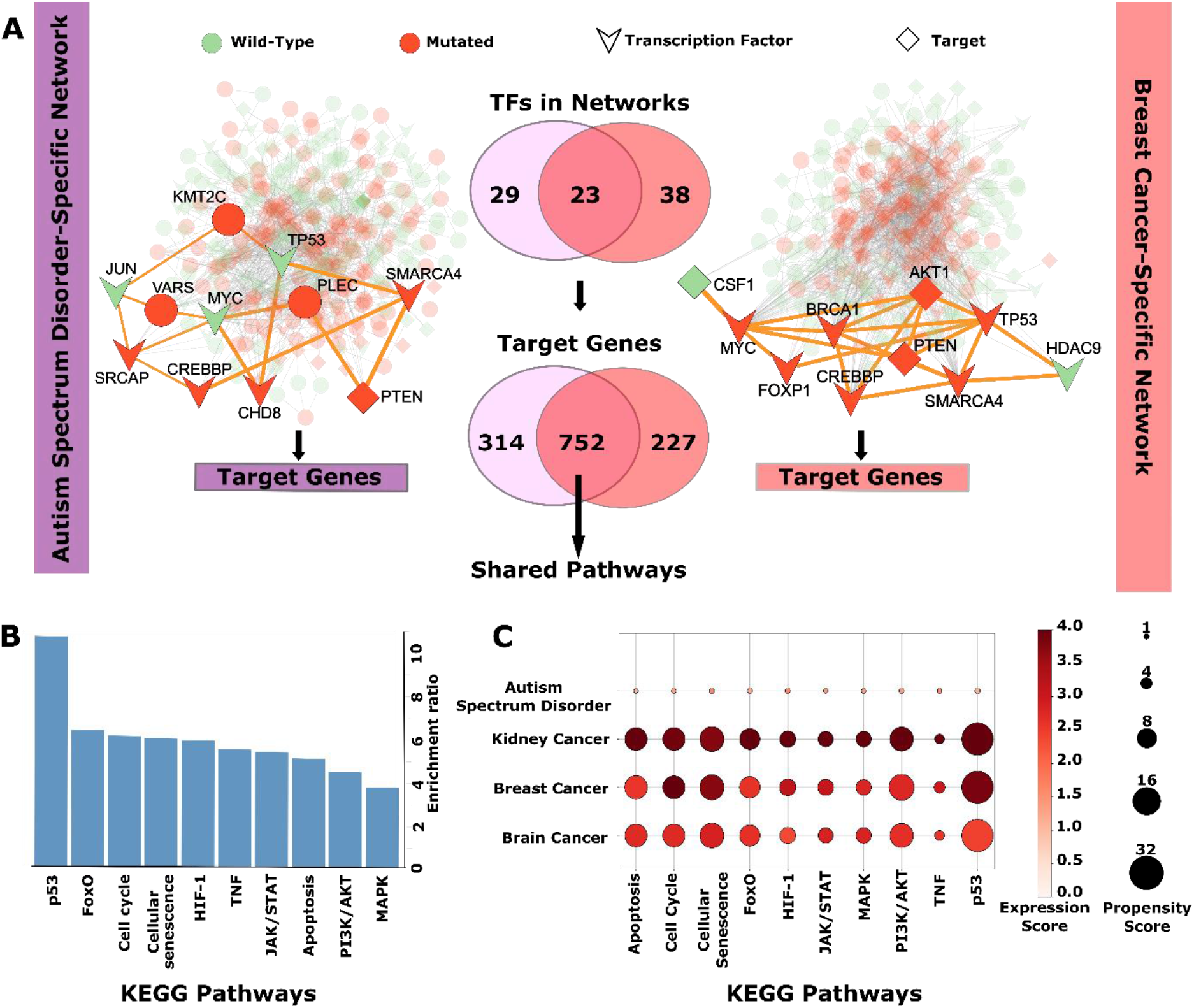
ASD- and breast cancer-specific networks regulating common pathways. **(A)** Disease-specific network reconstruction for ASD and breast cancer is performed by using pyPARAGON tool, where the frequently mutated genes are used as seeds. The nodes in reconstructed networks involve wild type (green circle), mutated genes (red circle), TFs (chevron), and TF-targets (diamond). The complete ASD-specific network (left side) features the mutated proteins (SRCAP, BRG1, PTEN, etc.) in ASD cases and reveals disease-associated proteins (Jun, p53, and Myc). The breast cancer-specific network (right side) illustrates driver genes, although some driver genes, such as *TP53* and *MYC*, are not frequently mutated in ASD. Both ASD- and breast cancer-specific networks involve 23 common TFs targeting 752 common genes. These common targets are employed to identify shared pathways. Abbreviations: BRG1, brahma-related gene 1, a.k.a. SMARCA4; SRCAP, SNF2-related CREBBP activator protein; CREBBP, cAMP response element binding protein; CHD8, chromodomain helicase DNA-binding protein 8; CSF1, macrophage colony-stimulating factor 1; HD9, histone deacetylase 9; FOXP1, forkhead box protein P1. **(B**) Overrepresentation analysis determines significant shared pathways (FDR ≤ 0.05) related to cell differentiation and proliferation among KEGG pathways. The pathways include MAPK, PI3K/AKT, and JAK/STAT. These shared TF-target genes play a significant role in cell fate by altering the signal strength and flow, as well as cell cycle and cellular senescence. Abbreviations: HIF-1, hypoxia-inducible factor 1; TNF, tumor necrosis factor. (**C**) Signal changes in shared pathways are illustrated with the expression scores of pathways, the mean of the absolute *z*-scores of proteins in a given pathway. We define expression scores as the mean of the absolute *z*-scores of proteins in a given pathway to indicate the magnitude of the deviation from the average expression values of the normal samples, regardless of the direction of the change. The vulnerability of common pathways to mutation is measured with a propensity score, the average unique mutation in the pathway. The darker red represents a higher change in expression scores of genes in the pathway, and the larger circle shows a higher mutation propensity for the corresponding pathway. ASD has the most minor signal differences and mutation propensities compared to all cancer types in shared pathways, where kidney cancer has the highest signal difference. However, there is an insignificant difference in mutation propensities amongst cancer types. The higher expression scores in cancer types point to stronger signal changes in pathways critical for cell fate, such as proliferation and differentiation. The higher propensity scores in cancer reveal that cancer mutations tend to group in shared pathways. Thus, shared pathways are more vulnerable to cancer than ones in ASD. However, mutation loads and signal deviations on the shared pathways might make ASD patients more fragile to cancer onset.

All TFs in ASD- and breast cancer-specific networks regulate 752 commonly targeted genes. The disease models in both networks can use different wiring mechanisms to control shared pathways since different TFs control the transcription of the same genes. Overrepresentation analysis of these common targets demonstrated that shared pathways, including p53, FOXO (forkhead box O), PI3K/AKT, MAPK, and JAK/STAT (Janus kinase/signal transducer and activator of transcription) signaling pathways, are regulated by different TFs (Figure 4B).

### Gene expression and signaling strength point to differentiation in ASD and proliferation in cancer

Following the construction of the networks and identification of the TFs and their targets, we focused on the signal levels in the constructed networks through an analysis based on differential gene expressions from healthy and disease samples (see Materials and Methods). We averaged the absolute values of the differential expression of pathway participants and defined them as the expression score of the given pathway to measure the signal change in these pathways. The expression scores of the overrepresented pathways demonstrated that ASD generated significantly lower signal strength than breast, brain, and kidney cancers (Figure 4C), influencing the cell cycle at the G1 phase. The change in stimulus and feedback loops regulate signaling intensity and duration (Mendoza et al., 2011). Overexpression and multiple mutation combinations on these pathways disrupt cellular processes and can govern disease development.

The expression profiles of ASD in shared pathways emphasize differentiation. Differentiation reduces the proliferative advantage for the cells and increases their resistance to oncogenic mutations (Demeter et al., 2022). Mutations in ASD are mostly embryonic; they do not accumulate over time as cancer mutations do. The propensity score of pathways, which demonstrates the probabilities of mutations on a gene in a pathway, reveals that mutations in cancer tend to accumulate in these pathways. Shared pathways in ASD do not have high propensity scores. The already existing mutational burden makes ASD patients more susceptible to multifactorial and/or polygenic diseases, like cancer (Nussinov et al., 2022d; Parenti et al., 2020; Rauen, 2013). At the same time, their weak/moderate effect can bring about cell cycle arrest and impact the differentiation capabilities of cells.

### TFs regulating common pathways underscore the trends of differentiation in NDDs and proliferation in cancer

For a more in-depth analysis, we compared 71 TFs regulating common pathways through the expression profiles of ASD and breast cancer patients. We observed that 21 TFs that have distinct expression profiles in ASD and cancer are clustered into three groups. Cluster-1 and Cluster-2 demonstrated a distinct separation, while Cluster-3 includes genes that do not show a clear difference in the heatmap of gene expressions (Figure 5A). The genes in Cluster-1, such as *MCM2*, *STAT1*, *BRCA1*, and *MCM5*, are overexpressed in the cancer samples. These genes mostly play a role in cell proliferation, and their overexpression in cancer promotes cell division and growth (Gong et al., 2019; Shimizu et al., 2012; Wu et al., 2018; Yousef et al., 2017). On the contrary, ASD samples have relatively lower expression levels for TFs that control cell proliferation. *STAT1* has dual roles in both differentiation and proliferation; it also acts as a tumor suppressor and an oncogene in cancer. The genes in Cluster-2, such as *JUN*, *SMAD3*, *SMAD4*, and *KLF2*, play a role in cell differentiation (Hou et al., 2018; Mariani et al., 2007; Yang et al., 2016; Yang and Jiang, 2020). Their moderate expression levels in ASD suggest that they can maintain the cell differentiation state. To reveal the signal flow starting from these TFs, we defined the regulatory interaction in common pathways by identifying target genes of these TFs. Since one TF can also target other TFs in the same pathway, we extended the regulatory interactions with targeted TFs and their targeted genes (Figure 5B). Expression profiles of differentiation and proliferation appear moderate in ASD, which suggests weak signal activation in cell proliferation (Nussinov et al., 2023). However, the suppression of differentiation and the overexpression of proliferation indicate strong activation of the proliferation state in cancer.

**Figure 5.**
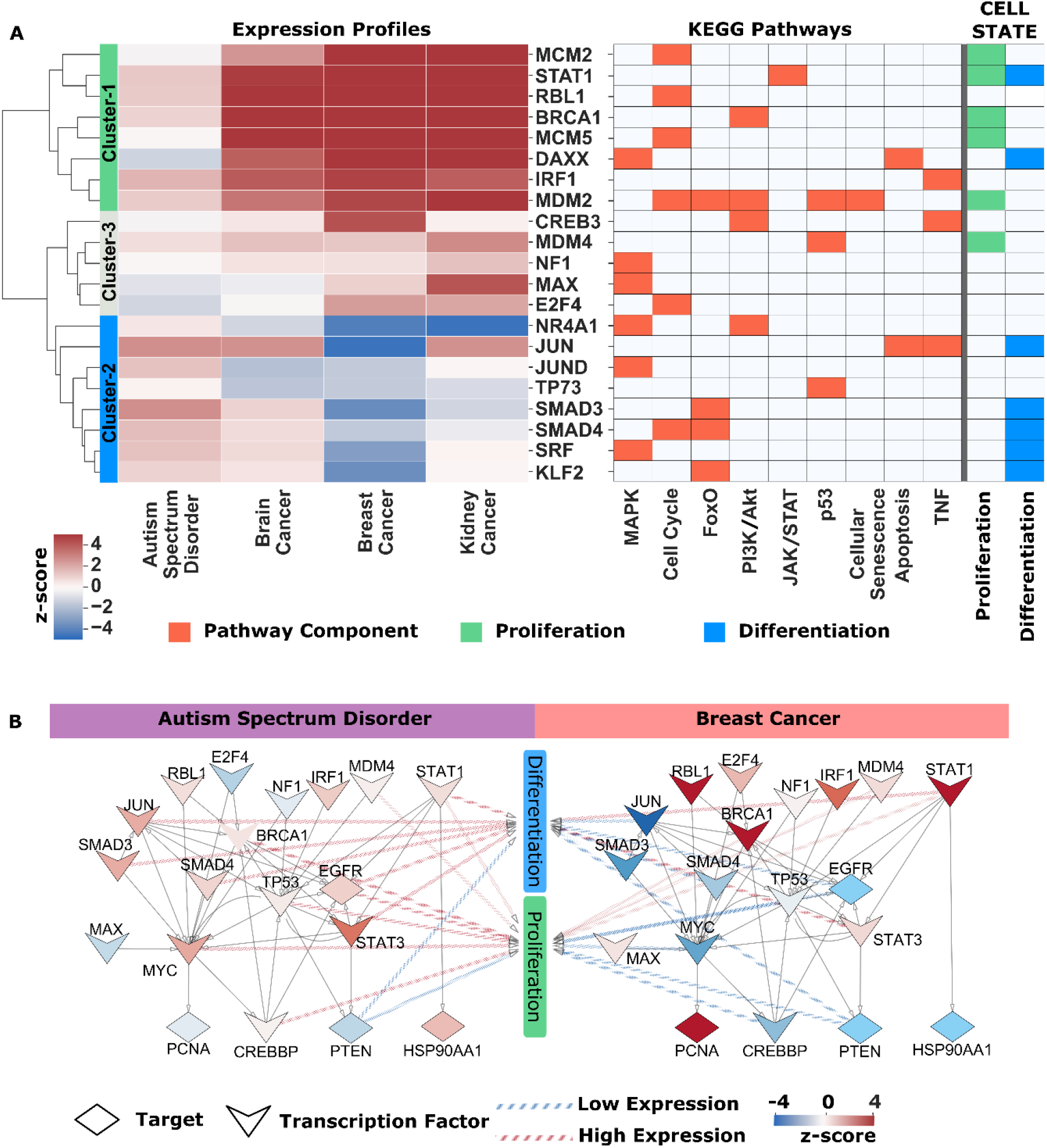
Differential expression profiles in shared pathways. **(A)** Differentiated expression profiles of TFs in shared pathways. There were 71 TFs in shared pathways that determine cell fate via changes in signal levels. However, 21 TFs were identified to be at least one time differentially expressed more (less) in ASD than in other cancer types. On the left hand, the heatmap of these differentially expressed genes (high in red, low in blue) clustered expression *z*-scores into three groups. On the right hand, the pathways TFs belong to, and related cell states (proliferation, green; differentiation, blue) are demonstrated. *MCM2*, *STAT1*, *BRCA1*, *MCM5*, *DAXX*, *IRF1*, and *MDM2* in cluster-1 are highly expressed in cancers, while *NR4A1*, *JUN*, *JUND*, *TP73*, *SMAD3*, *SMAD4*, *SRF*, and *KLF2* in cluster-2 are highly expressed in Autism. Genes more expressed in cancer types than in ASD mainly belong to the proliferation state, while genes related to differentiation are predominantly more expressed in ASD than in cancer types. (**B**) Differences between proliferation and differentiation on shared pathways. The signal flows from TFs (chevron) to targets (diamond) in common parts of ASD- and breast cancer-specific networks and in shared pathways were demonstrated with *z*-scores. The low and high expression levels were illustrated with blue to red, respectively. The relationship between cell state and proteins is represented with arrows whose color also demonstrates the level of expressions, low or high. Differentiation-related proteins, such as Jun, SMAD3, and SMAD4, mainly have low expression profiles in breast cancer, while most are highly expressed in ASD. PTEN, EGFR, and STAT1, related to proliferation and differentiation, have similar expression profiles. Abbreviations: E2F4, E2F transcription factor 4; RBL1, retinoblastoma-like protein 1; NF1, neurofibromin; IRF1, interferon regulatory factor 1; BRCA1, breast cancer type 1 susceptibility protein; SMAD, mothers against decapentaplegic; EGFR, epidermal growth factor receptor; PCNA, proliferating cell nuclear antigen; CREBBP, cAMP response element binding protein; Hsp90α, heat shock protein 90α.

## Discussion

### Moderate and strong escalation in signaling levels reflect the total number of activated molecules in NDDs and cancer, respectively

Here, we comprehensively analyzed mutations, transcriptomic data, and PPI networks of NDDs and cancer patients to comprehend why some mutations can promote cancer while others abet NDDs, and why the same mutations can support both phenotypes. We surmised that cancer mutations are connected to elevated signaling levels, while NDD mutations encode sustained but low levels. We further surmised that signaling levels are largely determined by the total number of molecules that the mutations activate, either alone or in combination, along with the cell type-specific expression levels of the mutant protein and other proteins in the relevant pathways, the timing of the emergence of the mutation (inherited or during embryonic development, or sporadic), as well as additional factors (Nussinov et al., 2022d). Ample data indicate that even high expression levels of an unmutated protein can already provoke cancer. Cancer involves uncontrolled cell proliferation, whereas NDDs are connected to anomalies in the development of the nervous system. Proliferation and differentiation take place in both cancer and NDDs. Since NDDs are mostly related to dysregulated differentiation, mutations in genes regulating chromatin organization rank high. Risk genes for NDDs include more than a third of the cancer driver genes, and NDDs and cancer share hallmarks of cell division and growth (Yaeger and Corcoran, 2019; Zhao and Luo, 2022), thus proliferation and differentiation (Nussinov et al., 2022d; Qi et al., 2016). In brain cells, embryonic mutations in both pathways give rise to NDDs (Borrie et al., 2017). Hundreds of genes are implicated in NDDs; however, they are involved in few conserved pathways regulating transcription, including chromatin accessibility, and synaptic signaling (Nussinov et al., 2022d; Parenti et al., 2020; Sahin and Sur, 2015). PI3K/mTOR and Ras/MAPK are frequently linked with synaptic dysregulation (Longo and Klann, 2021; Nussinov et al., 2022a, 2022d; Sahin and Sur, 2015). Proteins in the Wnt, BMP/TGF-β (bone morphogenetic protein/transforming growth factor-β), SHH (sonic hedgehog), FGF (fibroblast growth factor), and RA (retinoic acid) pathways, are also involved in autistic brain development (Kumar et al., 2019). Gene expression profiles of 22 cancer types and frontal cortical tissues from ASD patients identified similarities in genes and pathways (Forés-Martos et al., 2019).

### NDDs share phenotypic and clinical commonalities

The tumor suppressor phosphatase and tensin homolog (PTEN), which carries germline and *de novo* mutations in NDD patients, is related to cancer and several NDDs, collectively named PHTS. The NDDs include phenotypes such as Cowden syndrome (CS), Bannayan-Riley-Ruvalcaba syndrome (BRRS), Proteus syndrome (PS), Proteus-like syndrome (PSL), macrocephaly, and ASD. NDDs often overlap mutation-wise and genome-wise (Frazier et al., 2021; Orrico et al., 2009; Skelton et al., 2020). Among these, deletions, and duplications of the 16p11.2 region are common. About 48% of deletion carriers and 63% of duplication carriers have at least one psychiatric diagnosis (Niarchou et al., 2019; Walsh and Bracken, 2011). RASopathies, which include Noonan syndrome (NS), cardiofaciocutaneous (CFC) syndrome, neurofibromatosis type 1 (NF1), and Legius syndrome (LS), are NDDs that result from overactivation of the MAPK pathway due to germline mutations and/or overexpression in embryogenesis (Gross et al., 2020; Hebron et al., 2022; Nussinov et al., 2022d). Their phenotypic overlaps may emerge due to shared proteins/pathways as in the case of *PIK3CA*-related overgrowth spectrum (PROS), PS, and CS which share phenotypic characteristics with RASopathies (Simanshu et al., 2017). The commonality of cancer and RASopathies prompted MEK (MAPK kinase) inhibitors and Ras-targeted therapies for some RASopathies like selumetinib for NF1 patients (Andelfinger et al., 2019; Cox et al., 2015; Dombi et al., 2016; Hebron et al., 2022).

Although there is a strong association between PTEN germline mutations and cancer– PHTS–they have also been described in patients with ASDs (Cummings et al., 2022; Nussinov et al., 2022d; Skelton et al., 2020). PTEN mutations linked to ASD can lead to an unstable but still catalytically active gene product (Chang et al., 2008). C124S, G129R, H118P, H123Q, E157G, F241S, D252G, N276S, and D326N are autism-related; A39P, N48K, L108P, L112P, and R130L are PHTS-related mutations (Spinelli et al., 2015). AKT, downstream of PTEN, signaling was suppressed in all seven ASD-related PTEN mutations where PTEN was affected but functional. On the other hand, AKT phosphorylation was promoted by all five PTEN mutations in severe PHTS cases, suggesting that variants with partial loss of PTEN function are predominant in ASD patients (Spinelli et al., 2015). Thus, catalytically inactive PTEN mutant is connected to tumor phenotypes, partially active PTEN to ASD (Papa et al., 2014; Rodríguez-Escudero et al., 2011).

Dysregulation of the PI3K/AKT/mTOR pathway is a primary factor in NDDs, including megalencephaly (also known as “large brain”), microcephaly (sometimes known as “small brain”), ASD, intellectual disability, schizophrenia, and epilepsy (Wang et al., 2017). Mosaic gain-of-function mutations in the *PIK3CA* gene lead to PROS, with clinical outcomes such as excessive tissue growth, blood vessel abnormalities, and scoliosis (Crunkhorn, 2018; Venot et al., 2018). Among ∼200 individuals with *PIK3CA* mosaic mutations, highly activating hotspot mutations were associated with severe brain and/or body overgrowth, whilst fewer activating mutations were linked to more mild somatic overgrowth and mostly brain overgrowth (Dobyns and Mirzaa, 2019; Mirzaa et al., 2016). R88Q, V344M, and G914R mutations were identified in PI3Kα patients with macrocephaly and developmental delay or ASD (Yeung et al., 2017).

### Distinct rewired interactions in shared ASD and breast cancer pathways

We further pursued the complex relationship between genotype and phenotype by constructing disease-specific networks for ASD and breast cancer. We observed distinct PPIs in shared pathways controlling the cell cycle. These rewired interactions could be a reason why shared pathways have different signal strengths in ASD and cancer. Under physiological conditions, MAPK and PI3K/AKT/mTOR pathways coregulate the cell cycle through feedback loops to control cell functions, including growth and division. In cancers, they are frequently hyperactivated (Ersahin et al., 2015; Thorpe et al., 2014; Vanhaesebroeck et al., 2010). The PI3K/AKT pathway is also critical in early embryonic development and maintenance of stem cell pluripotency through inhibition of the MAPK proliferation pathway (Bi et al., 1999; Hall, 2004; Peng et al., 2003; Yu and Cui, 2016). The strength of the signaling perturbations induced by the mutations is manifested in weak/moderate and strong signaling changes, epitomized by ASD and breast cancer, respectively. Strong signals enhance proliferation, and weak/moderate signals may drive cell cycle exit in differentiation (Eastman et al., 2020).

### Differential interactions of cell cycle CDKs in NDDs and cancer, and late cancer detection outcome for individuals with NDD

TF complexes are primarily involved in cell cycle regulation through their targets, such as E2F mediating CDK that accelerates proliferation (DeGregori et al., 1997; Tadesse et al., 2019). In the breast cancer-specific network, CDK4 interacts with MAPK1, JAK3, and p53, promoting proliferation (Scheiblecker et al., 2020). In the ASD-specific network, TF complexes such as forkhead box protein G1 (FOXG1) and sex determining region Y-box 2 (SOX2), also implicated in microcephaly, play critical roles in lineage determination, neural stem/progenitor cell proliferation, and maintenance of pluripotency (Hou et al., 2020; Li et al., 2013). In NDDs, these TFs can promote premature senescence and dysregulated differentiation via distinct pathways such as Wnt and Hippo (Nussinov et al., 2016). In a study of the English population, half of the decedents with intellectual disabilities and cancer were at stage IV when diagnosed (Heslop et al., 2022), which suggested involvement of the canonical Wnt pathway during brain morphogenesis and non-canonical in cancer cell migration and metastasis (Corda and Sala, 2017). Cancer onset in NDDs can be undetected until stage IV since the slow cell division in the NDDs retards mutational accumulation (Heslop et al., 2022). Alternatively, we expect the early mutational burden will render NDD patients more vulnerable to cancer (Nussinov et al., 2022d; Parenti et al., 2020), with faster cancer progression and higher mortality. Where statistics are available, the mortality of cancer patients with intellectual disabilities was reported to be approximately 1.5 times higher than the general population (Cuypers et al., 2022). These results suggest that cancer initiation and progression differ in individuals with NDD than in the broad apparent NDD-free population, with different outcomes via common pathways.

### TF expression profiles differ in differentiating and proliferating cells

The expression scores of TFs were grouped based on proliferation and differentiation. TFs enhancing proliferation were mainly overexpressed in cancers while relatively low-expressed in ASD. Proliferating cells are more vulnerable to mutations than differentiating ones, both since dividing cells have less time to repair DNA damage than quiescent cells, and with more replication cycles, there is a higher chance for mutations (Bielas and Heddle, 2000; Demeter et al., 2022). As to TFs in the differentiation state, ASD has relatively higher expression profiles, while there are significantly low-expression profiles in cancers. In cancers, high expression couples with the accumulation of mutations, cell growth, and metastasis (Demeter et al., 2022).

Finally, immunity could be viewed as a common factor in NDDs and cancer (Nussinov et al., 2022a, 2022d). Multiple pathways related to immunity can be dysregulated in NDDs due to the coevolution of the immune and nervous systems (Nussinov et al., 2022a; Zengeler and Lukens, 2021). Signaling pathways related to immunity, such as Wnt, Notch, JAK/STAT, and Hippo, also play roles in cancer metastasis and drug resistance (Clara et al., 2020; Nussinov et al., 2016; Pisibon et al., 2021).

## Conclusions: Why then individuals with NDDs have a higher probability of cancer?

Our findings offer a mechanistic interpretation for *PTEN* and *PIK3CA* mutations frequently observed in cancer and NDD samples, which may form the basis for functional and detailed structural analysis, including molecular dynamics simulations (Jang et al., 2023). Comparing expression scores of shared pathways by leveraging the transcriptomic profiles of NDDs and cancer samples revealed that NDD samples have higher expression scores for genes functioning in differentiation than proliferation. These findings provide an essential step toward understanding the etiology of the two different pathologies, NDDs, and cancer. Despite having common signaling pathways, their regulation and differences in signal levels enhance different cell states: proliferation for cancer and differentiation for NDDs.

Comparisons of the time windows of NDDs and cancer frequently conclude that while cancer is predominantly caused by somatic mutations and alterations in signaling and transcriptional programs, NDDs are primarily linked to germline mutations that express during embryonic development. A recent study has similarly suggested that mutations in cancer susceptibility genes are not necessarily inherited or somatic; they can also arise throughout embryogenesis as a result of errors occurring during cell division (Pareja et al., 2022). These *mosaic mutations*, occurring in early embryogenesis, were suspected to be associated with some rare cancers. Genetic changes associated with RASopathies are believed to be often sporadic, not inherited. Along these lines, according to the NCI page (“NCI Dictionary of Cancer Terms,” 2011), this means that typically multiple family members do not share the same NDDs.

Different from this view, here our thesis is that inherited and *de novo* mutations (missense or truncation) can be major causes of NDDs such as intellectual disability, ASD, epilepsy (Brunet et al., 2021; Chau et al., 2021; Deciphering Developmental Disorders Study, 2017; Iossifov et al., 2014), and cancer (Nussinov et al., 2021, 2019a; Nussinov and Tsai, 2015). As in cancer (Nussinov et al., 2021), more than one mutation is required for observable symptomatic NDDs. Our premise is that family members can harbor these NDD germline mutations; however, they are not diagnosed as having the disorder. Their offsprings are, however, already susceptible to it. Individuals with NDDs have higher probabilities of eventually coming down with cancer (Liu et al., 2022); (Cuypers et al., 2022; Liu et al., 2022; Nordentoft et al., 2021), likely due to the preexistence of the mutations in the shared proteins, making them more susceptible. Patients with autism have an increased mutation load in genes that drive cancer (Darbro et al., 2016). We hypothesize that strong driver mutations in cell growth/division pathways are chiefly responsible for uncontrolled cell proliferation in cancer. NDDs’ weak/moderate strength mutations may be a reason why inherited NDDs have not been identified in a parent while predisposing an offspring to it. An additional mutation promotes NDD clinical manifestation. It may be inherited from the other parent or emerge during embryogenesis. It may also promote cancer by providing companion mutations.

Here, we employed de novo mutations in ∼10,000 samples with NDDs from denovo-db and somatic mutations in ∼10,000 tumor samples from TCGA. We observed that around 40% of the 19,439 mutant genes in TCGA are also altered in NDD samples. 1504 of the 14,133 distinct NDD mutations are present in TCGA. On the other hand, TCGA contains 1060 distinct driver mutations, whereas known cancer driver mutations and NDD only share 23 mutations. This result suggests that common mutations across the two pathologies do exist, although they are typically less potent than cancer drivers. Especially, PTEN and PI3Kα possess a range of mutations scattered through their protein sequences that are either common or disease specific. This work argues for the examination of such mutations even in undiagnosed family members and follows their combination in the offspring. It further supports the consideration of cancer pharmacology in NDD patients.

## Materials and methods

### Data collection and processing

NDD mutations were obtained from denovo-db (Turner et al., 2017) which holds a collection of human germlines *de novo* variants of 20 phenotypes including but not limited to ASD, and intellectual disability NDDs. Variants from two ASD studies were collected by targeted sequencing of different patients coming from two different studies, while the remaining datasets come from either whole exome or whole genome studies. The phenotypes, the number of samples, unique mutated genes and unique mutations are given in Figure 1B. We mapped the genomic coordinates to the proteins to obtain the amino acid changes on the protein level using VarMap (Stephenson et al., 2019). We only kept the point mutations that map to the canonical protein sequences. After these filtering steps, we obtained a total of 14,133 unique mutations on 7907 genes from 9737 samples.

### TCGA

Somatic missense, nonsense and frameshift cancer mutations were downloaded from TCGA. There are 9703 tumor samples from 33 different cancer types in the annotation file where corresponding protein changes are also present. In total, we have 1,626,715 unique mutations on 19,438 genes. 7837 of these genes are also mutated in the NDD dataset. 11,601 of them are only mutated in TCGA, while there are only 70 genes that are mutated solely in NDDs.

### Cancer drivers

A list of cancer driver mutations was downloaded from the Cancer Genome Interpreter (CGI) (Tamborero et al., 2018), which is available as the Catalog of Validated Oncogenic Mutations on their website. We only used missense or nonsense mutations, resulting in an analysis of 3688 driver mutations belonging to 237 genes.

### Expression datasets

We utilized processed RNA expression data from ASD, breast, kidney, and brain cancer samples (Table 1) (Forés-Martos et al., 2019). The ASD dataset was an integrated dataset from the frontal cortex samples in three studies and covered 34 ASD samples and 130 controls. We employed integrated datasets for breast, kidney, and brain cancers that are composed of 7, 10, and 8 studies, respectively. Differential gene expression meta-analyses scored 3579 genes in ASD and 11629 genes in cancer cohorts with *z*-scores.

**Table 1.** RNA expression datasets.

| Phenotype | Cases | Control | Datasets |
| --- | --- | --- | --- |
| ASD | 34 | 130 | GSE28475, GSE28521, and Gupta (Gupta et al., 2014) |
| Brain cancer | 942 | 104 | GSE4290, GSE9385, GSE74195, GSE68848, GSE15824, GSE42656, GSE44971, GSE50161 |
| Breast cancer | 1494 | 249 | GSE10810, GSE31448, GSE42568, GSE54002, GSE65216, GSE45827, GSE29431 |
| Kidney cancer | 400 | 266 | GSE11151, GSE77199, GSE47032, GSE53757, GSE53000, GSE66272, GSE68417, GSE71963, GSE40435, GSE7635 |

### Pathway and network analyses

#### Inference of disease-specific networks

ASD and breast cancer-specific networks were reconstructed with frequently mutated genes and known PPIs. In cases of observations seen in at least 3 patients, 190 genes were selected as seed nodes in ASD. 168 genes were retrieved from the Cancer Genome Interpreter (CGI) and recruited as the seed nodes of breast cancer (Tamborero et al., 2018). The reference network, HIPPIE v2.3, comprises 19437 proteins and 779301 PPIs (Alanis-Lobato et al., 2017). Each interaction in HIPPIE was scored with a confidence score that was computationally optimized and weighted by the amount and quality of the experimental evidence of PPI. The network inference tool, pyPARAGON (PAgeRAnk-flux on Graphlet-guided-network for multi-Omic data integratioN), inferred ASD and breast cancer-specific networks by scoring interactions with PageRank Flux and identifying validated graphlet motifs, the union of which constructs a graphlet-guided network (GGN) (Arici and Tuncbag, 2023). The PageRank algorithm weighted all nodes in a reference network. We then used the flux computation to weight the edges (Rubel and Ritz, 2020). Significant graphlet motifs with seed nodes in the reference established the GGN. Highly scored interactions in GGN were assembled in our disease-specific networks. We used PARAGON with the following parameters: α = 0.5, where α is the probability of walking to neighbor nodes, τ = 0.8, where τ is a scaling factor to select a set of top-ranked edges from GGN. This algorithm stops adding edges when the number of interactions reaches 2000.

#### Identification of common pathways

TFs and their targets, retrieved from TRRUST v2, were parsed in disease-specific networks, and TFs in these networks were called specific transcription factors (STF) (Han et al., 2018). The targets of STF were selected as regulated genes by disease-specific networks. These commonly regulated genes among ASD and breast cancer were utilized for overrepresentation analysis on WebGestalt to uncover the common pathways (*p* < 0.05 and FDR < 0.05) using manually curated open-source pathway databases, KEGG and Reactome (Gillespie et al., 2022; Kanehisa et al., 2022; Liao et al., 2019).

#### Pathway assessment metrics

The signal strength and mutation vulnerability of the pathways were evaluated. The expression level of each gene contributes to the signal deviation in the respective pathway. However, it is challenging to determine how this signal deviation affects the pathway because it contributes to multiple molecular functionalities. To measure the expression score (*ES*) of a given pathway, we calculated the average absolute signal differences of a pathway (Hwang, 2012; Kim et al., 2008; Levine et al., 2006) by applying the equation,

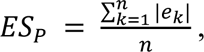

where *P* = (*G*, *E*, *U*), a pathway composed of genes/proteins (*g_1_*, *g_2_*, …, *g_n_*, ∋ *G*), expression of genes (|*e_1_*|, |*e_2_*|, …, |*e_n_|* ∋ *E*), and the number of unique mutations belonging to genes (*u_1_*, *u_2_*, …, *u_n_* ∋ *U*). We assessed the mutation vulnerability of a pathway by calculating the propensity score (*PS*) of a given pathway considering the number of unique mutations by using the equation,

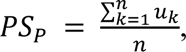

where the total number of individual mutations in the pathway was normalized with the number of gene members in the pathway.

### Visualization of mutations in protein sequences and 3D structures

We utilized the ProteinPaint tool (Zhou et al., 2016) to show NDD and cancer mutations on PTEN and PI3Kα. To map the mutations to the 3D structures of PTEN (PDB: 1D5R (Lee et al., 1999)) and PI3Kα (PDB: 4OVV (Miller et al., 2014)) we used PyMol.

## Acknowledgements

This project has been funded in whole or in part with federal funds from the National Cancer Institute, National Institutes of Health, under contract HHSN261201500003I. The content of this publication does not necessarily reflect the views or policies of the Department of Health and Human Services, nor does mention of trade names, commercial products, or organizations imply endorsement by the U.S. Government. This research was supported [in part] by the Intramural Research Program of the NIH, National Cancer Institute, Center for Cancer Research. NT has received support from the 2247-A National Outstanding Researchers Program of TUBITAK under the project number 121C292.

The results shown here are in part based upon data generated by the TCGA Research Network: https://www.cancer.gov/tcga. The results shown on NDD mutations are based on denovo-db, Seattle, WA (URL: <u>denovo-db.gs.washington.edu</u>).

## Competing Interests

The authors declare no competing interests.

## Authors’ contributions

Conceptualization: RN, NT, CJT

Data curation: BRY, MKA, HCD

Formal analysis: BRY, MKA, HCD

Methodology: BRY, MKA, HCD, CJT, HJ, RN, NT

Project administration: NT

Supervision: NT

Visualization: BRY, MKA, HCD, HJ

Writing – original draft: BRY, MKA, HCD, CJT, HJ, RN, NT

## Notes

### Competing Interest Statement

The authors have declared no competing interest.

## References

Abrahams BS, Arking DE, Campbell DB, Mefford HC, Morrow EM, Weiss LA, Menashe I, Wadkins T, Banerjee-Basu S, Packer A. 2013. SFARI Gene 2.0: a community-driven knowledgebase for the autism spectrum disorders (ASDs). Mol Autism 4:1–3.

Alanis-Lobato G, Andrade-Navarro MA, Schaefer MH. 2017. HIPPIE v2.0: enhancing meaningfulness and reliability of protein–protein interaction networks. Nucleic Acids Research. doi:10.1093/nar/gkw985

Andelfinger G, Marquis C, Raboisson M-J, Théoret Y, Waldmüller S, Wiegand G, Gelb BD, Zenker M, Delrue M-A, Hofbeck M. 2019. Hypertrophic Cardiomyopathy in Noonan Syndrome Treated by MEK-Inhibition. J Am Coll Cardiol 73:2237–2239.

Arici MK, Tuncbag N. 2023. pyPARAGON: PAgeRAnk-flux on Graphlet-guided-network for multi-Omic data integratioN. Zenodo. doi:10.5281/ZENODO.7736975

Bielas JH, Heddle JA. 2000. Proliferation is necessary for both repair and mutation in transgenic mouse cells. Proc Natl Acad Sci U S A 97:11391–11396.

Bi L, Okabe I, Bernard DJ, Wynshaw-Boris A, Nussbaum RL. 1999. Proliferative Defect and Embryonic Lethality in Mice Homozygous for a Deletion in the p110α Subunit of Phosphoinositide 3-Kinase *. J Biol Chem 274:10963–10968.

Borrie SC, Brems H, Legius E, Bagni C. 2017. Cognitive Dysfunctions in Intellectual Disabilities: The Contributions of the Ras-MAPK and PI3K-AKT-mTOR Pathways. Annu Rev Genomics Hum Genet 18:115–142.

Bragin E, Chatzimichali EA, Wright CF, Hurles ME, Firth HV, Bevan AP, Swaminathan GJ. 2014. DECIPHER: database for the interpretation of phenotype-linked plausibly pathogenic sequence and copy-number variation. Nucleic Acids Res 42:D993–D1000.

Brunet T, Jech R, Brugger M, Kovacs R, Alhaddad B, Leszinski G, Riedhammer KM, Westphal DS, Mahle I, Mayerhanser K, Skorvanek M, Weber S, Graf E, Berutti R, Necpál J, Havránková P, Pavelekova P, Hempel M, Kotzaeridou U, Hoffmann GF, Leiz S, Makowski C, Roser T, Schroeder SA, Steinfeld R, Strobl-Wildemann G, Hoefele J, Borggraefe I, Distelmaier F, Strom TM, Winkelmann J, Meitinger T, Zech M, Wagner M. 2021. De novo variants in neurodevelopmental disorders-experiences from a tertiary care center. Clin Genet 100:14–28.

Chang C-J, Mulholland DJ, Valamehr B, Mosessian S, Sellers WR, Wu H. 2008. PTEN nuclear localization is regulated by oxidative stress and mediates p53-dependent tumor suppression. Mol Cell Biol 28:3281–3289.

Chau KK, Zhang P, Urresti J, Amar M, Pramod AB, Chen J, Thomas A, Corominas R, Lin GN, Iakoucheva LM. 2021. Full-length isoform transcriptome of the developing human brain provides further insights into autism. Cell Rep 36:109631.

Chen M-H, Tsai S-J, Su T-P, Li C-T, Lin W-C, Cheng C-M, Chen T-J, Bai Y-M. 2022. Cancer risk in patients with bipolar disorder and unaffected siblings of such patients: A nationwide population-based study. Int J Cancer 150:1579–1586.

Chiang H-L, Liu C-J, Hu Y-W, Chen S-C, Hu L-Y, Shen C-C, Yeh C-M, Chen T-J, Gau SS-F. 2015. Risk of cancer in children, adolescents, and young adults with autistic disorder. J Pediatr 166:418–23.e1.

Clara JA, Monge C, Yang Y, Takebe N. 2020. Targeting signalling pathways and the immune microenvironment of cancer stem cells - a clinical update. Nat Rev Clin Oncol 17:204–232.

Corda G, Sala A. 2017. Non-canonical WNT/PCP signalling in cancer: Fzd6 takes centre stage. Oncogenesis 6:e364.

Cox AD, Der CJ, Philips MR. 2015. Targeting RAS Membrane Association: Back to the Future for Anti-RAS Drug Discovery? Clinical Cancer Research. doi:10.1158/1078-0432.ccr-14-3214

Crunkhorn S. 2018. Genetic disorders: PI3K inhibitor reverses overgrowth syndrome. Nat Rev Drug Discov 17:545.

Cummings K, Watkins A, Jones C, Dias R, Welham A. 2022. Behavioural and psychological features of PTEN mutations: a systematic review of the literature and meta-analysis of the prevalence of autism spectrum disorder characteristics. J Neurodev Disord 14:1.

Cuypers M, Schalk BWM, Boonman AJN, Naaldenberg J, Leusink GL. 2022. Cancer-related mortality among people with intellectual disabilities: A nationwide population-based cohort study. Cancer 128:1267–1274.

Darbro BW, Singh R, Zimmerman MB, Mahajan VB, Bassuk AG. 2016. Autism Linked to Increased Oncogene Mutations but Decreased Cancer Rate. PLoS One 11:e0149041.

Deciphering Developmental Disorders Study. 2017. Prevalence and architecture of de novo mutations in developmental disorders. Nature 542:433–438.

DeGregori J, Leone G, Miron A, Jakoi L, Nevins JR. 1997. Distinct roles for E2F proteins in cell growth control and apoptosis. Proc Natl Acad Sci U S A 94:7245–7250.

Demeter M, Derényi I, Szöllősi GJ. 2022. Trade-off between reducing mutational accumulation and increasing commitment to differentiation determines tissue organization. Nat Commun 13:1–10.

Dobyns WB, Mirzaa GM. 2019. Megalencephaly syndromes associated with mutations of core components of the PI3K-AKT-MTOR pathway: PIK3CA, PIK3R2, AKT3, and MTOR. Am J Med Genet C Semin Med Genet 181:582–590.

Dombi E, Baldwin A, Marcus LJ, Fisher MJ, Weiss B, Kim A, Whitcomb P, Martin S, Aschbacher-Smith LE, Rizvi TA, Wu J, Ershler R, Wolters P, Therrien J, Glod J, Belasco JB, Schorry E, Brofferio A, Starosta AJ, Gillespie A, Doyle AL, Ratner N, Widemann BC. 2016. Activity of Selumetinib in Neurofibromatosis Type 1-Related Plexiform Neurofibromas. N Engl J Med 375:2550–2560.

Eastman AE, Chen X, Hu X, Hartman AA, Pearlman Morales AM, Yang C, Lu J, Kueh HY, Guo S. 2020. Resolving Cell Cycle Speed in One Snapshot with a Live-Cell Fluorescent Reporter. Cell Rep 31:107804.

Ersahin T, Tuncbag N, Cetin-Atalay R. 2015. The PI3K/AKT/mTOR interactive pathway. Mol Biosyst 11:1946–1954.

Forés-Martos J, Catalá-López F, Sánchez-Valle J, Ibáñez K, Tejero H, Palma-Gudiel H, Climent J, Pancaldi V, Fañanás L, Arango C, Parellada M, Baudot A, Vogt D, Rubenstein JL, Valencia A, Tabarés-Seisdedos R. 2019. Transcriptomic metaanalyses of autistic brains reveals shared gene expression and biological pathway abnormalities with cancer. Mol Autism 10:17.

Frazier TW, Jaini R, Busch RM, Wolf M, Sadler T, Klaas P, Hardan AY, Martinez-Agosto JA, Sahin M, Eng C, Warfield SK, Scherrer B, Dies K, Filip-Dhima R, Gulsrud A, Hanson E, Phillips JM, the Developmental Synaptopathies Consortium. 2021. Cross-level analysis of molecular and neurobehavioral function in a prospective series of patients with germline heterozygous PTEN mutations with and without autism. Molecular Autism. doi:10.1186/s13229-020-00406-6

Garces de Los Fayos Alonso I, Liang H-C, Turner SD, Lagger S, Merkel O, Kenner L. 2018. The Role of Activator Protein-1 (AP-1) Family Members in CD30-Positive Lymphomas. Cancers 10. doi:10.3390/cancers10040093

García-Gutiérrez L, Bretones G, Molina E, Arechaga I, Symonds C, Acosta JC, Blanco R, Fernández A, Alonso L, Sicinski P, Barbacid M, Santamaría D, León J. 2019. Myc stimulates cell cycle progression through the activation of Cdk1 and phosphorylation of p27. Sci Rep 9:18693.

Gillespie M, Jassal B, Stephan R, Milacic M, Rothfels K, Senff-Ribeiro A, Griss J, Sevilla C, Matthews L, Gong C, Deng C, Varusai T, Ragueneau E, Haider Y, May B, Shamovsky V, Weiser J, Brunson T, Sanati N, Beckman L, Shao X, Fabregat A, Sidiropoulos K, Murillo J, Viteri G, Cook J, Shorser S, Bader G, Demir E, Sander C, Haw R, Wu G, Stein L, Hermjakob H, D’Eustachio P. 2022. The reactome pathway knowledgebase 2022. Nucleic Acids Res 50:D687–D692.

Gong B, Ma M, Yang X, Xie W, Luo Y, Sun T. 2019. MCM5 promotes tumour proliferation and correlates with the progression and prognosis of renal cell carcinoma. Int Urol Nephrol 51:1517–1526.

Gross AM, Frone M, Gripp KW, Gelb BD, Schoyer L, Schill L, Stronach B, Biesecker LG, Esposito D, Hernandez ER, Legius E, Loh ML, Martin S, Morrison DK, Rauen KA, Wolters PL, Zand D, McCormick F, Savage SA, Stewart DR, Widemann BC, Yohe ME. 2020. Advancing RAS/RASopathy therapies: An NCI-sponsored intramural and extramural collaboration for the study of RASopathies. American Journal of Medical Genetics Part A. doi:10.1002/ajmg.a.61485

Gupta S, Ellis SE, Ashar FN, Moes A, Bader JS, Zhan J, West AB, Arking DE. 2014. Transcriptome analysis reveals dysregulation of innate immune response genes and neuronal activity-dependent genes in autism. Nat Commun 5:5748.

Hall M. 2004. Faculty Opinions recommendation of mTOR is essential for growth and proliferation in early mouse embryos and embryonic stem cells. Faculty Opinions – Post-Publication Peer Review of the Biomedical Literature. doi:10.3410/f.1020418.232572

Han H, Cho J-W, Lee S, Yun A, Kim H, Bae D, Yang S, Kim CY, Lee M, Kim E, Lee S, Kang B, Jeong D, Kim Y, Jeon H-N, Jung H, Nam S, Chung M, Kim J-H, Lee I. 2018. TRRUST v2: an expanded reference database of human and mouse transcriptional regulatory interactions. Nucleic Acids Res 46:D380–D386.

Hebron KE, Hernandez ER, Yohe ME. 2022. The RASopathies: from pathogenetics to therapeutics. Dis Model Mech 15. doi:10.1242/dmm.049107

Heslop P, Cook A, Sullivan B, Calkin R, Pollard J, Byrne V. 2022. Cancer in deceased adults with intellectual disabilities: English population-based study using linked data from three sources. BMJ Open 12:e056974.

Hou P-S, hAilín DÓ, Vogel T, Hanashima C. 2020. Transcription and Beyond: Delineating FOXG1 Function in Cortical Development and Disorders. Front Cell Neurosci 14:35.

Hou Z, Wang Z, Tao Y, Bai J, Yu B, Shen J, Sun H, Xiao L, Xu Y, Zhou J, Wang Z, Geng D. 2018. KLF2 regulates osteoblast differentiation by targeting of Runx2. Lab Invest 99:271–280.

Huang K-L, Mashl RJ, Wu Y, Ritter DI, Wang J, Oh C, Paczkowska M, Reynolds S, Wyczalkowski MA, Oak N, Scott AD, Krassowski M, Cherniack AD, Houlahan KE, Jayasinghe R, Wang L-B, Zhou DC, Liu D, Cao S, Kim YW, Koire A, McMichael JF, Hucthagowder V, Kim T-B, Hahn A, Wang C, McLellan MD, Al-Mulla F, Johnson KJ, Cancer Genome Atlas Research Network, Lichtarge O, Boutros PC, Raphael B, Lazar AJ, Zhang W, Wendl MC, Govindan R, Jain S, Wheeler D, Kulkarni S, Dipersio JF, Reimand J, Meric-Bernstam F, Chen K, Shmulevich I, Plon SE, Chen F, Ding L. 2018. Pathogenic Germline Variants in 10,389 Adult Cancers. Cell 173:355–370.e14.

Hwang S. 2012. Comparison and evaluation of pathway-level aggregation methods of gene expression data. BMC Genomics 13 Suppl 7:S26.

Iossifov I, O’Roak BJ, Sanders SJ, Ronemus M, Krumm N, Levy D, Stessman HA, Witherspoon KT, Vives L, Patterson KE, Smith JD, Paeper B, Nickerson DA, Dea J, Dong S, Gonzalez LE, Mandell JD, Mane SM, Murtha MT, Sullivan CA, Walker MF, Waqar Z, Wei L, Willsey AJ, Yamrom B, Lee Y-H, Grabowska E, Dalkic E, Wang Z, Marks S, Andrews P, Leotta A, Kendall J, Hakker I, Rosenbaum J, Ma B, Rodgers L, Troge J, Narzisi G, Yoon S, Schatz MC, Ye K, McCombie WR, Shendure J, Eichler EE, State MW, Wigler M. 2014. The contribution of de novo coding mutations to autism spectrum disorder. Nature 515:216–221.

Jang H, Chen J, Iakoucheva LM, Nussinov R. 2023. How PTEN mutations degrade function at the membrane and life expectancy of carriers of mutations in the human brain. bioRxiv. doi:10.1101/2023.01.26.525746

Jang H, Smith IN, Eng C, Nussinov R. 2021. The mechanism of full activation of tumor suppressor PTEN at the phosphoinositide-enriched membrane. iScience 24:102438.

Jiang C-C, Lin L-S, Long S, Ke X-Y, Fukunaga K, Lu Y-M, Han F. 2022. Signalling pathways in autism spectrum disorder: mechanisms and therapeutic implications. Signal Transduct Target Ther 7:229.

Kanehisa M, Furumichi M, Sato Y, Kawashima M, Ishiguro-Watanabe M. 2022. KEGG for taxonomy-based analysis of pathways and genomes. Nucleic Acids Res. doi:10.1093/nar/gkac963

Kao H-T, Buka SL, Kelsey KT, Gruber DF, Porton B. 2010. The correlation between rates of cancer and autism: an exploratory ecological investigation. PLoS One 5:e9372.

Kim T-M, Yim S-H, Jeong Y-B, Jung Y-C, Chung Y-J. 2008. PathCluster: a framework for gene set-based hierarchical clustering. Bioinformatics 24:1957–1958.

Koire A, Katsonis P, Kim YW, Buchovecky C, Wilson SJ, Lichtarge O. 2021. A method to delineate de novo missense variants across pathways prioritizes genes linked to autism. Sci Transl Med 13. doi:10.1126/scitranslmed.abc1739

Kumar S, Reynolds K, Ji Y, Gu R, Rai S, Zhou CJ. 2019. Impaired neurodevelopmental pathways in autism spectrum disorder: a review of signaling mechanisms and crosstalk. J Neurodev Disord 11:10.

Lee JO, Yang H, Georgescu MM, Di Cristofano A, Maehama T, Shi Y, Dixon JE, Pandolfi P, Pavletich NP. 1999. Crystal structure of the PTEN tumor suppressor: implications for its phosphoinositide phosphatase activity and membrane association. Cell 99:323–334.

Levine DM, Haynor DR, Castle JC, Stepaniants SB, Pellegrini M, Mao M, Johnson JM. 2006. Pathway and gene-set activation measurement from mRNA expression data: the tissue distribution of human pathways. Genome Biol 7:R93.

Liao Y, Wang J, Jaehnig EJ, Shi Z, Zhang B. 2019. WebGestalt 2019: gene set analysis toolkit with revamped UIs and APIs. Nucleic Acids Res 47:W199–W205.

Li B, Li K, Tian D, Zhou Q, Xie Y, Fang Z, Wang X, Luo T, Wang Z, Zhang Y, Wang Y, Chen Q, Meng Q, Zhao G, Li J. 2020. De novo mutation of cancer-related genes associates with particular neurodevelopmental disorders. J Mol Med 98:1701–1712.

Li C, Ito H, Fujita K, Shiwaku H, Qi Y, Tagawa K, Tamura T, Okazawa H. 2013. Sox2 transcriptionally regulates PQBP1, an intellectual disability-microcephaly causative gene, in neural stem progenitor cells. PLoS One 8:e68627.

Liu M, Liu X, Suo P, Gong Y, Qu B, Peng X, Xiao W, Li Y, Chen Y, Zeng Z, Lu Y, Huang T, Zhao Y, Liu M, Li L, Chen Y, Zhou Y, Liu G, Yao J, Chen S, Song L. 2020. The contribution of hereditary cancer-related germline mutations to lung cancer susceptibility. Transl Lung Cancer Res 9:646–658.

Liu Q, Yin W, Meijsen JJ, Reichenberg A, Gådin JR, Schork AJ, Adami H-O, Kolevzon A, Sandin S, Fang F. 2022. Cancer risk in individuals with autism spectrum disorder. Ann Oncol 33:713–719.

Longo F, Klann E. 2021. Reciprocal control of translation and transcription in autism spectrum disorder. EMBO Rep 22:e52110.

Mariani O, Brennetot C, Coindre J-M, Gruel N, Ganem C, Delattre O, Stern M-H, Aurias A. 2007. JUN oncogene amplification and overexpression block adipocytic differentiation in highly aggressive sarcomas. Cancer Cell 11:361–374.

Mendoza MC, Er EE, Blenis J. 2011. The Ras-ERK and PI3K-mTOR pathways: cross-talk and compensation. Trends Biochem Sci 36:320–328.

Mighell TL, Evans-Dutson S, O’Roak BJ. 2018. A Saturation Mutagenesis Approach to Understanding PTEN Lipid Phosphatase Activity and Genotype-Phenotype Relationships. Am J Hum Genet 102:943–955.

Milenković T, Pržulj N. 2008. Uncovering Biological Network Function via Graphlet Degree Signatures. Cancer Informatics. doi:10.4137/cin.s680

Miller MS, Schmidt-Kittler O, Bolduc DM, Brower ET, Chaves-Moreira D, Allaire M, Kinzler KW, Jennings IG, Thompson PE, Cole PA, Mario Amzel L, Vogelstein B, Gabelli SB. 2014. Structural basis of nSH2 regulation and lipid binding in PI3Kα. Oncotarget. doi:10.18632/oncotarget.2263

Mirzaa G, Timms AE, Conti V, Boyle EA, Girisha KM, Martin B, Kircher M, Olds C, Juusola J, Collins S, Park K, Carter M, Glass I, Krägeloh-Mann I, Chitayat D, Parikh AS, Bradshaw R, Torti E, Braddock S, Burke L, Ghedia S, Stephan M, Stewart F, Prasad C, Napier M, Saitta S, Straussberg R, Gabbett M, O’Connor BC, Keegan CE, Yin LJ, Lai AHM, Martin N, McKinnon M, Addor M-C, Boccuto L, Schwartz CE, Lanoel A, Conway RL, Devriendt K, Tatton-Brown K, Pierpont ME, Painter M, Worgan L, Reggin J, Hennekam R, Tsuchiya K, Pritchard CC, Aracena M, Gripp KW, Cordisco M, Van Esch H, Garavelli L, Curry C, Goriely A, Kayserilli H, Shendure J, Graham J, Guerrini R, Dobyns WB. 2016. PIK3CA-associated developmental disorders exhibit distinct classes of mutations with variable expression and tissue distribution. JCI Insight. doi:10.1172/jci.insight.87623

Muiños F, Martínez-Jiménez F, Pich O, Gonzalez-Perez A, Lopez-Bigas N. 2021. In silico saturation mutagenesis of cancer genes. Nature 596:428–432.

NCI Dictionary of Cancer Terms. 2011.. National Cancer Institute. https://www.cancer.gov/publications/dictionaries/cancer-terms

Niarchou M, Chawner SJRA, Doherty JL, Maillard AM, Jacquemont S, Chung WK, Green-Snyder L, Bernier RA, Goin-Kochel RP, Hanson E, Linden DEJ, Linden SC, Raymond FL, Skuse D, Hall J, Owen MJ, van den Bree MBM. 2019. Psychiatric disorders in children with 16p11.2 deletion and duplication. Transl Psychiatry 9:8.

Nordentoft M, Plana-Ripoll O, Laursen TM. 2021. Cancer and schizophrenia. Curr Opin Psychiatry 34:260–265.

Nussinov R, Jang H, Tsai C-J, Cheng F. 2019a. Precision medicine review: rare driver mutations and their biophysical classification. Biophys Rev 11:5–19.

Nussinov R, Tsai C-J. 2015. “Latent drivers” expand the cancer mutational landscape. Curr Opin Struct Biol 32:25–32.

Nussinov R, Tsai C-J, Jang H. 2022a. Neurodevelopmental disorders, immunity, and cancer are connected. iScience 25. doi:10.1016/j.isci.2022.104492

Nussinov R, Tsai C-J, Jang H. 2022b. Allostery, and how to define and measure signal transduction. Biophys Chem 283:106766.

Nussinov R, Tsai C-J, Jang H. 2022c. A New View of Activating Mutations in Cancer. Cancer Res 82:4114–4123.

Nussinov R, Tsai C-J, Jang H. 2022d. How can same-gene mutations promote both cancer and developmental disorders? Sci Adv 8:eabm2059.

Nussinov R, Tsai C-J, Jang H. 2021. Anticancer drug resistance: An update and perspective. Drug Resist Updat 59:100796.

Nussinov R, Tsai C-J, Jang H. 2019b. Why Are Some Driver Mutations Rare? Trends Pharmacol Sci 40:919–929.

Nussinov R, Tsai C-J, Jang H, Korcsmáros T, Csermely P. 2016. Oncogenic KRAS signaling and YAP1/β-catenin: Similar cell cycle control in tumor initiation. Semin Cell Dev Biol 58:79–85.

Nussinov R, Yavuz BR, Arici MK, Demirel HC, Zhang M, Liu Y, Tsai C-J, Jang H, Tuncbag N. 2023. Neurodevelopmental disorders, like cancer, are connected to impaired chromatin remodelers, PI3K/mTOR and PAK1-regulated MAPK. Biophys Rev. doi:DOI: 10.1007/s12551-023-01054-9

Orrico A, Galli L, Buoni S, Orsi A, Vonella G, Sorrentino V. 2009. Novel PTEN mutations in neurodevelopmental disorders and macrocephaly. Clin Genet 75:195–198.

Papa A, Wan L, Bonora M, Salmena L, Song MS, Hobbs RM, Lunardi A, Webster K, Ng C, Newton RH, Knoblauch N, Guarnerio J, Ito K, Turka LA, Beck AH, Pinton P, Bronson RT, Wei W, Pandolfi PP. 2014. Cancer-Associated PTEN Mutants Act in a Dominant-Negative Manner to Suppress PTEN Protein Function. Cell. doi:10.1016/j.cell.2014.03.027

Pareja F, Ptashkin RN, Brown DN, Derakhshan F, Selenica P, da Silva EM, Gazzo AM, Da Cruz Paula A, Breen K, Shen R, Marra A, Zehir A, Benayed R, Berger MF, Ceyhan-Birsoy O, Jairam S, Sheehan M, Patel U, Kemel Y, Casanova-Murphy J, Schwartz CJ, Vahdatinia M, Comen E, Borsu L, Pei X, Riaz N, Abramson DH, Weigelt B, Walsh MF, Hadjantonakis A-K, Ladanyi M, Offit K, Stadler ZK, Robson ME, Reis-Filho JS, Mandelker D. 2022. Cancer-Causative Mutations Occurring in Early Embryogenesis. Cancer Discov 12:949–957.

Parenti I, Rabaneda LG, Schoen H, Novarino G. 2020. Neurodevelopmental Disorders: From Genetics to Functional Pathways. Trends in Neurosciences. doi:10.1016/j.tins.2020.05.004

Pejaver V, Urresti J, Lugo-Martinez J, Pagel KA, Lin GN, Nam H-J, Mort M, Cooper DN, Sebat J, Iakoucheva LM, Mooney SD, Radivojac P. 2020. Inferring the molecular and phenotypic impact of amino acid variants with MutPred2. Nat Commun 11:5918.

Peng X-D, Xu P-Z, Chen M-L, Hahn-Windgassen A, Skeen J, Jacobs J, Sundararajan D, Chen WS, Crawford SE, Coleman KG, Hay N. 2003. Dwarfism, impaired skin development, skeletal muscle atrophy, delayed bone development, and impeded adipogenesis in mice lacking Akt1 and Akt2. Genes Dev 17:1352–1365.

Piñero J, Queralt-Rosinach N, Bravo À, Deu-Pons J, Bauer-Mehren A, Baron M, Sanz F, Furlong LI. 2015. DisGeNET: a discovery platform for the dynamical exploration of human diseases and their genes. Database 2015:bav028.

Pisibon C, Ouertani A, Bertolotto C, Ballotti R, Cheli Y. 2021. Immune Checkpoints in Cancers: From Signaling to the Clinic. Cancers 13:4573.

Portelli S, Barr L, de Sá AGC, Pires DEV, Ascher DB. 2021. Distinguishing between PTEN clinical phenotypes through mutation analysis. Comput Struct Biotechnol J 19:3097– 3109.

Qi H, Dong C, Chung WK, Wang K, Shen Y. 2016. Deep Genetic Connection Between Cancer and Developmental Disorders. Hum Mutat 37:1042–1050.

Qing T, Mohsen H, Marczyk M, Ye Y, O’Meara T, Zhao H, Townsend JP, Gerstein M, Hatzis C, Kluger Y, Pusztai L. 2020. Germline variant burden in cancer genes correlates with age at diagnosis and somatic mutation burden. Nat Commun 11:2438.

Rashed WM, Marcotte EL, Spector LG. 2022. Germline Mutations as a Cause of Childhood Cancer. JCO Precis Oncol 6:e2100505.

Rauen KA. 2013. The RASopathies. Annu Rev Genomics Hum Genet 14:355–369.

Rodríguez-Escudero I, Oliver MD, Andrés-Pons A, Molina M, Cid VJ, Pulido R. 2011. A comprehensive functional analysis of PTEN mutations: implications in tumor- and autism-related syndromes. Hum Mol Genet 20:4132–4142.

Rubel T, Ritz A. 2020. Augmenting signaling pathway reconstructions. bioRxiv. doi:10.1101/2020.06.16.155853

Sahin M, Sur M. 2015. Genes, circuits, and precision therapies for autism and related neurodevelopmental disorders. Science 350. doi:10.1126/science.aab3897

Scheiblecker L, Kollmann K, Sexl V. 2020. CDK4/6 and MAPK-Crosstalk as Opportunity for Cancer Treatment. Pharmaceuticals 13. doi:10.3390/ph13120418

Shimizu Y, Luk H, Horio D, Miron P, Griswold M, Iglehart D, Hernandez B, Killeen J, ElShamy WM. 2012. BRCA1-IRIS Overexpression Promotes Formation of Aggressive Breast Cancers. PLoS One 7:e34102.

Simanshu DK, Nissley DV, McCormick F. 2017. RAS Proteins and Their Regulators in Human Disease. Cell. doi:10.1016/j.cell.2017.06.009

Singh G, Driever PH, Sander JW. 2005. Cancer risk in people with epilepsy: the role of antiepileptic drugs. Brain 128:7–17.

Singh G, Fletcher O, Bell GS, McLean AE, Sander JW. 2009. Cancer mortality amongst people with epilepsy: a study of two cohorts with severe and presumed milder epilepsy. Epilepsy Res 83:190–197.

Skelton PD, Stan RV, Luikart BW. 2020. The Role of PTEN in Neurodevelopment. Mol Neuropsychiatry 5:60–71.

Spinelli L, Black FM, Berg JN, Eickholt BJ, Leslie NR. 2015. Functionally distinct groups of inherited PTEN mutations in autism and tumour syndromes. J Med Genet 52:128–134.

Stephenson JD, Laskowski RA, Nightingale A, Hurles ME, Thornton JM. 2019. VarMap: a web tool for mapping genomic coordinates to protein sequence and structure and retrieving protein structural annotations. Bioinformatics 35:4854–4856.

Stratton M. 2008. Patterns of somatic mutation in human cancer genomes. European Journal of Cancer Supplements. doi:10.1016/s1359-6349(08)71197-2

Tadesse S, Caldon EC, Tilley W, Wang S. 2019. Cyclin-Dependent Kinase 2 Inhibitors in Cancer Therapy: An Update. J Med Chem 62:4233–4251.

Tamborero D, Rubio-Perez C, Deu-Pons J, Schroeder MP, Vivancos A, Rovira A, Tusquets I, Albanell J, Rodon J, Tabernero J, de Torres C, Dienstmann R, Gonzalez-Perez A, Lopez-Bigas N. 2018. Cancer Genome Interpreter annotates the biological and clinical relevance of tumor alterations. Genome Med 10:25.

Thorpe LM, Yuzugullu H, Zhao JJ. 2014. PI3K in cancer: divergent roles of isoforms, modes of activation and therapeutic targeting. Nat Rev Cancer 15:7–24.

Turner TN, Yi Q, Krumm N, Huddleston J, Hoekzema K, F Stessman HA, Doebley A-L, Bernier RA, Nickerson DA, Eichler EE. 2017. denovo-db: a compendium of human de novo variants. Nucleic Acids Res 45:D804–D811.

Vanhaesebroeck B, Guillermet-Guibert J, Graupera M, Bilanges B. 2010. The emerging mechanisms of isoform-specific PI3K signalling. Nat Rev Mol Cell Biol 11:329–341.

Venot Q, Blanc T, Rabia SH, Berteloot L, Ladraa S, Duong J-P, Blanc E, Johnson SC, Hoguin C, Boccara O, Sarnacki S, Boddaert N, Pannier S, Martinez F, Magassa S, Yamaguchi J, Knebelmann B, Merville P, Grenier N, Joly D, Cormier-Daire V, Michot C, Bole-Feysot C, Picard A, Soupre V, Lyonnet S, Sadoine J, Slimani L, Chaussain C, Laroche-Raynaud C, Guibaud L, Broissand C, Amiel J, Legendre C, Terzi F, Canaud G. 2018. Targeted therapy in patients with PIK3CA-related overgrowth syndrome. Nature 558:540–546.

Walsh KM, Bracken MB. 2011. Copy number variation in the dosage-sensitive 16p11.2 interval accounts for only a small proportion of autism incidence: a systematic review and meta-analysis. Genet Med 13:377–384.

Wang L, Zhou K, Fu Z, Yu D, Huang H, Zang X, Mo X. 2017. Brain Development and Akt Signaling: the Crossroads of Signaling Pathway and Neurodevelopmental Diseases. J Mol Neurosci 61:379–384.

Wong CW, Or PMY, Wang Y, Li L, Li J, Yan M, Cao Y, Luk HM, Tong TMF, Leslie NR, Lo IF-M, Choy KW, Chan AML. 2018. Identification of a PTEN mutation with reduced protein stability, phosphatase activity, and nuclear localization in Hong Kong patients with autistic features, neurodevelopmental delays, and macrocephaly. Autism Research. doi:10.1002/aur.1950

Wu W, Wang X, Shan C, Li Y, Li F. 2018. Minichromosome maintenance protein 2 correlates with the malignant status and regulates proliferation and cell cycle in lung squamous cell carcinoma. OTT 11:5025–5034.

Xu X, Zhou Y, Feng X, Li X, Asad M, Li D, Liao B, Li J, Cui Q, Wang E. 2020. Germline genomic patterns are associated with cancer risk, oncogenic pathways, and clinical outcomes. Sci Adv 6. doi:10.1126/sciadv.aba4905

Yaeger R, Corcoran RB. 2019. Targeting Alterations in the RAF-MEK Pathway. Cancer Discov 9:329–341.

Yang J, Jiang W. 2020. The Role of SMAD2/3 in Human Embryonic Stem Cells. Front Cell Dev Biol 0. doi:10.3389/fcell.2020.00653

Yang J, Wang J, Zeng Z, Qiao L, Zhuang L, Jiang L, Wei J, Ma Q, Wu M, Ye S, Gao Q, Ma D, Huang X. 2016. Smad4 is required for the development of cardiac and skeletal muscle in zebrafish. Differentiation 92:161–168.

Yeung KS, Tso WWY, Ip JJK, Mak CCY, Leung GKC, Tsang MHY, Ying D, Pei SLC, Lee SL, Yang W, Chung BH-Y. 2017. Identification of mutations in the PI3K-AKT-mTOR signalling pathway in patients with macrocephaly and developmental delay and/or autism. Mol Autism 8:1–11.

Yousef EM, Furrer D, Laperriere DL, Tahir MR, Mader S, Diorio C, Gaboury LA. 2017. MCM2: An alternative to Ki-67 for measuring breast cancer cell proliferation. Mod Pathol 30:682–697.

Yu JSL, Cui W. 2016. Proliferation, survival and metabolism: the role of PI3K/AKT/mTOR signalling in pluripotency and cell fate determination. Development 143:3050–3060.

Zengeler KE, Lukens JR. 2021. Innate immunity at the crossroads of healthy brain maturation and neurodevelopmental disorders. Nat Rev Immunol 21:454–468.

Zhang M, Jang H, Nussinov R. 2019. The mechanism of PI3Kα activation at the atomic level. Chem Sci 10:3671–3680.

Zhao J, Luo Z. 2022. Discovery of Raf Family Is a Milestone in Deciphering the Ras-Mediated Intracellular Signaling Pathway. Int J Mol Sci 23. doi:10.3390/ijms23095158

Zhou X, Edmonson MN, Wilkinson MR, Patel A, Wu G, Liu Y, Li Y, Zhang Z, Rusch MC, Parker M, Becksfort J, Downing JR, Zhang J. 2016. Exploring genomic alteration in pediatric cancer using ProteinPaint. Nat Genet 48:4–6.

